# Pathways regulating the excitability of sympathetic neurons derived from human induced pluripotent stem cells

**DOI:** 10.64898/2026.05.23.727423

**Authors:** Laura Fedele, Margot Maurer, Andrew Tinker, David A. Andersson

## Abstract

Postganglionic sympathetic neurons not only regulate target organs but also their own intrinsic activity and can be further modulated by parasympathetic neurons. This crosstalk and automodulatory signalling have been implicated in cardiovascular disorders, with the majority of earlier work employing rodent models. Here, we have used human sympathetic neurons derived from induced pluripotent stem cells (hiPSCs), as a scalable human *in vitro* system with the aim to investigate these pathways. Efficient differentiation of hiPSCs into sympathetic neurons was confirmed using molecular characterisation for the expression of *PHOX2B*, *DBH*, *TH*, *PRPH*. We employed Ca^2+^ imaging and whole-cell patch-clamp electrophysiology, to examine the neuronal functional properties and found that hiPSC-derived sympathetic neurons recapitulate key physiological features of the rodent native counterparts. Most cells responded to nicotine and expressed functional α2 adrenergic and muscarinic receptors, involved in sympathetic autoregulation and parasympathetic crosstalk. We further demonstrated that α2 adrenergic and muscarinic receptors inhibit membrane excitability (increased rheobase, hyperpolarisation, reduced input resistance) and that both types of receptors converge on inwardly rectifying K^+^ channels (GIRK) as effectors. The GIRK blocker Tertiapin-Q significantly reduced the α2 adrenergic and muscarinic responses, while the activator ML297 mimicked their action. Analysis of mouse stellate scRNA-seq confirmed that the receptors and GIRK subtypes studied here are prominently expressed in native sympathetic neurons. Overall, our data show that GIRK channels play an important role in the regulation of sympathetic neurons excitability and that hiPSC-derived neurons provide an attractive *in vitro* tool for drug discovery to study sympathetic autoregulation and parasympathetic-sympathetic crosstalk.

**KEY POINTS:**

- hiPSC-sympathetic neurons recapitulate key cellular pathways of native counterparts
- They express relevant receptors and ion channels involved in the inhibitory autoregulatory feedback and parasympathetic-sympathetic crosstalk
- Using a combination of RT-qPCR and functional recordings we identified inwardly rectifying K^+^ channels as downstream effectors of both α2 adrenergic and M2 muscarinic receptors. We cross-validated our findings with a mouse transcriptomic dataset from thoracic sympathetic ganglia.
- Overall, our data suggest that hiPSC-sympathetic neurons can be employed as a human *in vitro* platform to study cellular pathways and for drug discovery purposes.

Graphic abstract.
Summary of proposed modulation of sympathetic neuron activity via α2ARs and muscarinic receptors via GIRK channels
Functional recordings showed that muscarinic receptors and α2 adrenergic receptors reduce neuronal excitability, this effect was mimicked by activation of inwardly rectifying K^+^ channels (GIRK). Blockade of GIRK abolished the effects of both types of receptors, demonstrating that they converge on GIRK as a common effector. The muscarinic response was blocked by pertussis toxin, indicating an involvement of Gα_i/o_ downstream of the receptor, consistent with the high expression of *CHRM2* by RT-qPCR.

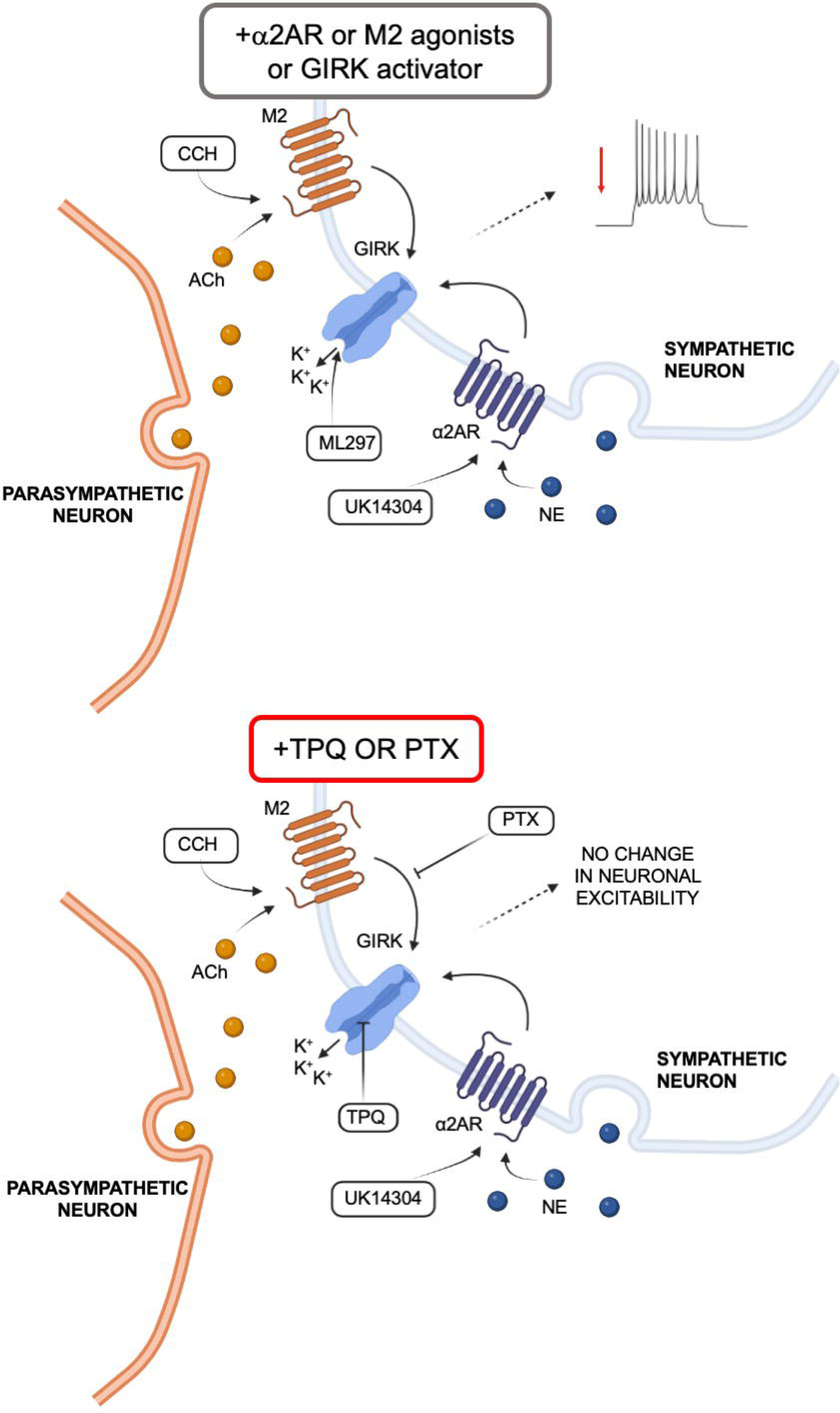

## INTRODUCTION

Sympathetic neurons employ a two-neuron chain system whereby the first neuron (preganglionic) resides in the central nervous system (spinal cord) and the postganglionic in the paravertebral chain (Wang *et al*., 2025). Postganglionic sympathetic neurons are characterised by the expression of the autonomic markers (PHOX2B) and genes involved in the synthesis pathway of their main neurotransmitter, norepinephrine: tyrosine hydroxylase (TH) and dopamine-β-hydroxylase (DBH) (Bos *et al*., 2025); they also release other co-transmitters (e.g. Neuropeptide Y, NPY)(Lundberg *et al*., 1983). Postganglionic sympathetic neurons innervate the target organs modulating their activity, for example they increase heart rate, contraction and conduction velocity (Fedele & Brand, 2020; Habecker *et al*., 2024; Herring *et al*., 2024). Sympathetic overactivity is associated with various cardiovascular conditions (e.g. catecholaminergic polymorphic tachycardia CPVT, long QT syndrome 1 LQTS1, hypertension, heart failure) (Habecker *et al*., 2024). Beta blockers, which act on β adrenergic receptors (the main targets of norepinephrine in cardiomyocytes), are among the most common therapeutics employed in the clinics to control cardiac activity; cardiac sympathetic denervation is part of the international guidelines for the treatment of CPVT and LQTS1(Herring *et al*., 2024). Beyond modulating target organs, sympathetic neurotransmitters can modulate the intrinsic activity of the neurons that release them through autoreceptors (Gilsbach & Hein, 2012). One example is the α2 adrenergic receptor (α2AR), which is located in prejunctional terminals where it regulates sympathetic neurotransmitter release (Lipscombe *et al*., 1989; Trendelenburg *et al*., 2001; Gilsbach & Hein, 2012). Consistent with a regulatory mechanism, α2ARs (α_2A and_ α_2A/C_) knockout mice present an increase in norepinephrine release (Hein *et al*., 1999), with their role also confirmed with pharmacological approaches (e.g. the α2ARs agonist UK14303) (Koh & Hille, 1997). Furthermore, α2ARs knockout mice present increased sympathetic tone, evident by higher baseline heart rate resulting in reduced cardiac function and abnormalities in cardiomyocytes structure (Brum *et al*., 2002). These mice were more susceptible to heart failure and cardiac hypertrophy (Gilsbach *et al*., 2010), which is consistent with the reduced function of α2ARs reported in heart failure patients (Aggarwal *et al*., 2001). Finally, α2ARs agonists were found to reduce mean firing rates (using multielectrode arrays) in hyperactive human induced pluripotent stem cells sympathetic neurons from familial dysautonomia patients (Wu *et al*., 2022).

Various reports have identified that the activity of sympathetic neurons is further regulated by parasympathetic neurons, with activation of muscarinic M2 receptors reducing neurotransmitter release (norepinephrine, neuropeptide Y) (Koh & Hille, 1997; Jani *et al*., 2025). Vagal tone plays a key role in counteracting sympathetic overactivity, with muscarinic receptors contributing to this mechanism (Jani *et al*., 2025). Autonomic imbalance has been reported in various cardiac disorders, including atrial fibrillation (Vandenberk *et al*., 2024) and heart failure (Van Weperen *et al*., 2023). In heart failure, sympathetic overactivity has been paired with reduced central and peripheral parasympathetic tone (Van Weperen *et al*., 2023), with the intracardiac ganglia (postganglionic neurons, mainly parasympathetic) presenting markers of degeneration (Van Weperen *et al*., 2023). Muscarinic receptors have long been studied for their role in atrial rhythm, but little work has been undertaken on their involvement in sympathetic neurons in physiological or pathophysiological states. Dissecting parasympathetic-sympathetic crosstalk in humans is difficult with the majority of published work focused on rodent models. Human models including induced pluripotent stem cell models have been proposed as a translational platform to dissect human neurophysiology (Luo *et al*., 2022).

The aim of our study was to explore cellular signalling pathways that could be targeted to modulate sympathetic neurons activity, employing a human system (hiPSC-derived sympathetic neurons). We explored the pathways of adrenergic and muscarinic receptors, since they are the receptors for the main neurotransmitters of the sympathetic and parasympathetic systems and are central for autoregulation and parasympathetic-sympathetic crosstalk.

We were guided by the pioneering work on adrenergic and muscarinic receptors in rodent superior cervical ganglia combined with overexpression studies that proposed a role of inwardly rectifying K^+^ channels (GIRK), N-type Ca^2+^ channels downstream both receptors types, and M-current (Lipscombe *et al*., 1989; Delmas *et al*., 1998; Shapiro *et al*., 2001; Fernández-Fernández *et al*., 2001). However, they did not investigate the overall effect of these receptors on sympathetic excitability nor the downstream effectors involved in their overall effect. To our knowledge no one attempted to dissect these pathways in a human system in sympathetic neurons, which would be important to identify potential drug targets and develop novel therapeutics.

We assessed the differentiated hiPSC-sympathetic neurons using RT-qPCR, immunocytochemistry for their autonomic and sympathetic lineage, and employed Ca^2+^ imaging and electrophysiological recordings to assess their function. Although hiPSC-derived sympathetic neurons have been generated by various groups (Winbo *et al*., 2020; Wu *et al*., 2024; Li *et al*., 2026), we provide the first quantification of the proportion of differentiated excitable cells responsive to nicotine. We further demonstrate that they express functional α2 adrenergic and muscarinic receptors, investigate their role in modulating sympathetic activity and we identified a key ion channel involved in this modulation employing established pharmacological tools. Our findings were consistent with the expressions of these receptors and the downstream ion channel genes in mouse sympathetic thoracic ganglia, scRNAseq dataset from (Furlan *et al*., 2016). Overall, our work suggests that hiPSC-sympathetic neurons can be employed for mechanistic exploration of at least these two modulatory pathways, but also for drug discovery purposes to exploit their high scalability and human relevance.

## MATERIAL AND METHODS

### Generation of Sympathetic Neurons from human induced pluripotent stem cells

The human induced pluripotent stem cell line HPSI0714ikute_4 (Kute4) (female line from a healthy donor) was employed in this study. This line was obtained through the Human Induced Pluripotent Stem Cell (HIPSCI) Initiative at King’s College London. Cells were maintained as explained in details in (Thomas *et al*., 2025). In brief, they were grown in vitronectin coated 6-well plates, using Stemflex media (Stemflex basal media, 10% Stemflex supplement and 1% of antimycotic, antibiotic), and passaged when 70% confluent. Before starting a differentiation, cells were seeded on Geltrex (90μg/ml in Stemflex basal media) coated 6-well plates and allowed to expand up to 70% confluence using Stemflex media. To generate hiPSC-derived sympathetic neurons, we adapted the protocol from Winbo et al 2020. We introduced a “neuronal enrichment” step, treating cells with 2μM cytosine beta-D-arabinofuranoside (AraC) at Day 21 or Day 24 for 72h, to inhibit the proliferation of non-neuronal mitotic cells. The neuronal enrichment step has been previously employed by us using AraC for hiPSC-derived sensory neurons with the same hiPSC cell line (Li *et al*., 2025*b*), the Zeltner group employed a similar approach but with Aphidicolin using a different protocol of hiPSC-derived sympathetic neurons (Wu *et al*., 2024). The composition of the “Neuronal media” was the following: Neurobasal, 1% N2, 2% B27, 1% Glutamax, 1% antibiotic/antimycotic, 0.2mM ascorbic acid; 0.2nM dbcAMP; 10ng/mL NGF; 10ng/mL BDNF; 10 ng/mL GDNF. Media, small molecules and supplements for each day were employed as the following. On Day 0-3 we employed Stemflex media, supplemented: on Day 0-1 with 500nM LDN and 10µM SB431542; on Day 2 with 500nM LDN, 10µM SB431542, 3µM CHIR99021, 10µM DAPT, 0.2µM PD173074. On Day 4-5 the media composition was 75% Stemflex and 25% Neuronal media supplemented with 3µM CHIR99021, 10µM DAPT, 0.2µM PD173074, 60ng/mL Shh C25II, 1µM purmorphamine. On Day 6-7 50% Stemflex and 50% Neuronal media, supplemented on Day 6 with 3µM CHIR99021, 10µM DAPT, 0.2µM PD173074, 60ng/mL Shh C25II, 1µM purmorphamine and on Day 7 with 60ng/mL Shh C25II, 1µM purmorphamine. Day 8 and 9 media was changed to 75% Neuronal media and 25% Stemflex supplemented with 60ng/mL Shh C25II and 1µM purmorphamine, Day 10 and 11 with Neuronal media (100%) with 60ng/mL Shh C25II and 1µM purmorphamine and 10ng/ml BMP4. Neuronal media was employed thereafter. On Day 12, cells were washed twice with PBS (0CaCl_2_, 0MgCl_2_) dissociated with TrypLE (7 minutes at 37°C), counted plated at a density of 50,000 cells on Geltrex coated (90μg/ml in Neurobasal media) coverslip (13mm). On both Day 12 and Day 13 neuronal media was supplemented with 10ng/mL BMP4 and on Day 12 10µM of Rock inhibitor (Y-27632) was also added to enhance the viability of dissociated cells. From Day 14 onwards, media was standard neuronal media. Media was changed every day for the first 14 days (day 14 included), on Day 17, 19, 21 and twice a week thereafter. On Day 21, neuronal media was supplemented with AraC, and on Day 17 and on Day 38 with Geltrex (1/250) to enhance cell adherence on coverslips. Neurons were employed at day 40-77 throughout the study.

### RNA extraction, cDNA and RT-qPCR

Three coverslips of neurons per batch or one well of a 6-well plate of undifferentiated hiPSC were considered one sample, respectively. Samples were collected using RLT buffer supplemented with β -mercaptoethanol (1% v/v). RNA was extracted using the Qiagen RNeasy Microkit (#74034) and quantified with Qubit^TM^ High Sensitivity RNA kit (#Q32852). cDNA synthesis was undertaken using Superscript III Reverse Transcriptase (#18080044). Real Time PCR (RT-PCR) was performed using the LightCycler 480 SYBR Green I Master Mix (#04707516001). Primers employed in this study are summarized in Table 1. Results were quantified as 2^-^ β^CT^ normalized to the housekeeping gene GAPDH. Five batches of neurons (D46-54) and 5 samples of corresponding undifferentiated iPSC (passage 28-40) were employed for the RT-qPCR, each dot represents the data from a sample from a distinct batch. The D46-54 timepoint was chosen because (Winbo *et al*., 2020) confirmed a mature-like profile at this timepoint, which was further confirmed by our functional electrophysiological data, including action potential characteristics, response to nicotine and to α2ARs and M2 agonists.

**Table 1.**
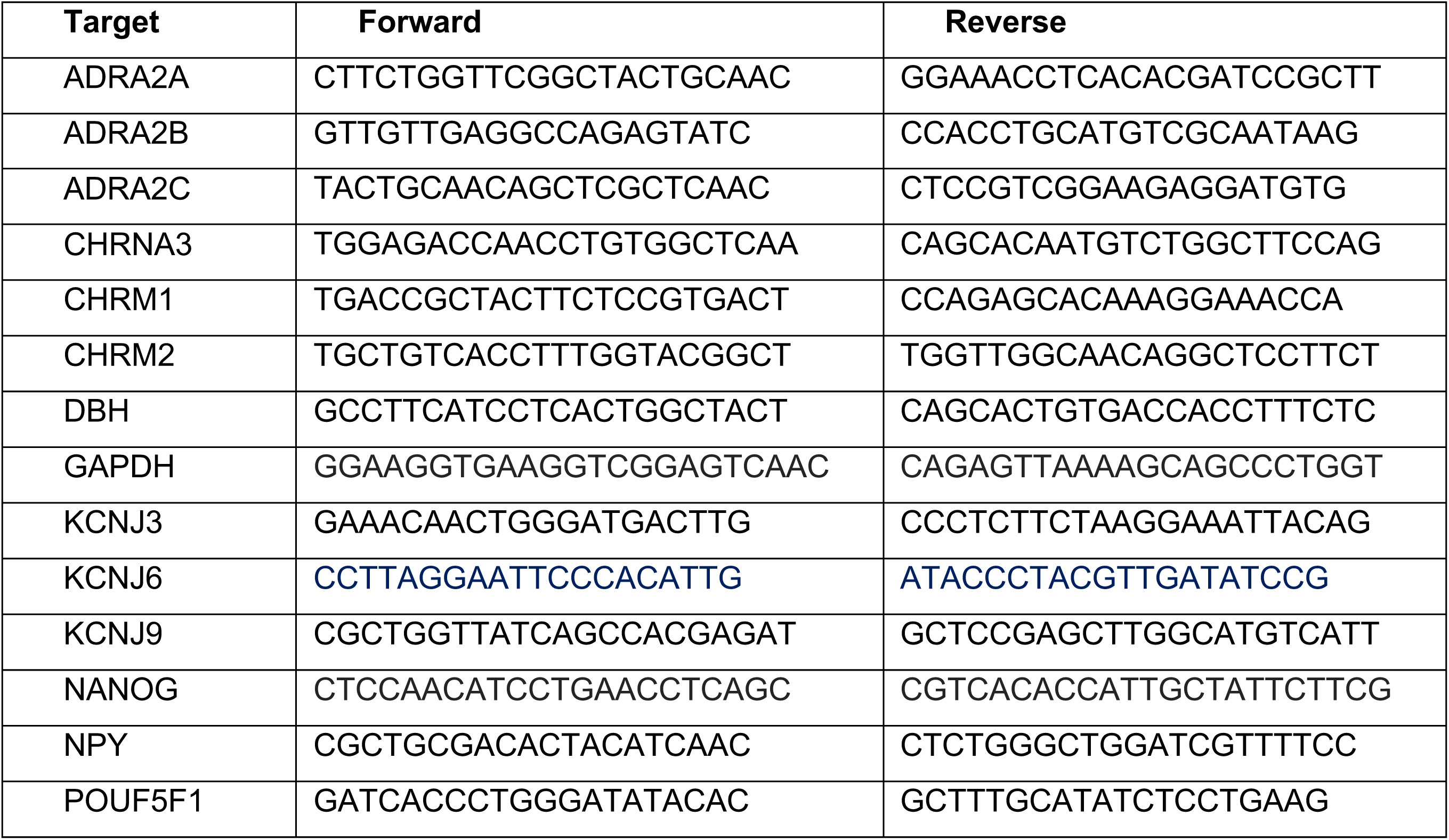
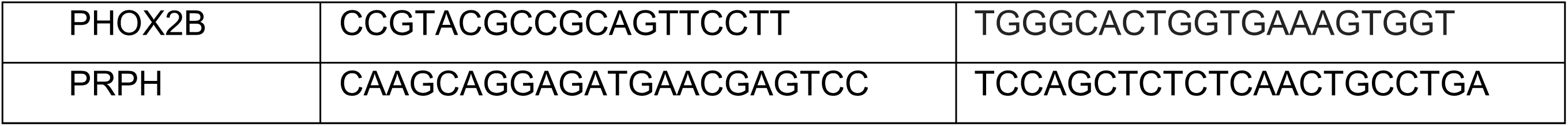
Primers used for qPCR.

**Table 2.** Primary antibodies.

| Primary antibody | Supplier | Dilution | Catalogue number |
| --- | --- | --- | --- |
| GIRK1 | Genetex | 1:500 | GTX108745 |
| Tyrosine Hydroxylase | Millipore | 1:500 | ab1542 |
| Phox2b | SantaCruz | 1:100 | sc-376997 |
| PGP9.5 | Abcam | 1:500 | ab108986 |
| PGP9.5 | Cedarlane | 1:2000 | CL7756AP |
| Beta III tubulin | Promega | 1:1000 | G7212 |

**Table 3.** Secondary Antibodies.

| Secondary antibody | Supplier | Dilution | Catalogue Number |
| --- | --- | --- | --- |
| Donkey Anti-rabbit 647 | Thermofisher | 1:1000 | A32795 |
| Donkey Anti-sheep 546 | Thermofisher | 1:1000 | A21098 |
| Donkey Anti-mouse 488 | Thermofisher | 1:1000 | A21202 |

### Immunocytochemistry and analyses

Cells were fixed with 2% PFA for 10 min at 37°C, washed three times with PBS and stored at 4°C until staining (up to 2 weeks). They were then permeabilised for 1 hour in blocking solution (0.2% Triton X-100 in PBS with 10% donkey serum). They were incubated in blocking solution containing primary antibody (or blocking solution without primary antibody for negative controls) overnight at room temperature. Afterwards, cells were washed three times with PBS and incubated for 1-2 hours in blocking solution containing the secondary antibodies. They were then washed with PBS three times and mounted on glass slides using DAPI-containing mounting media (DAPI-FluoromontG, #0100-20). Slides were left to dry overnight at room temperature and then stored at +4°C until imaging.

Coverslips were imaged using a Ti-E 2 AXR confocal microscope (Nikon Imaging Centre, King’s College London) with a 20x objective. Quantification of colocalization of DAPI positive nuclei with the autonomic marker PHOX2B or sympathetic marker TH was performed employing our previously published FIJI Macro (Zebochin *et al*., 2024), as explained in detail in (Li *et al*., 2025*b*).

### Whole cell patch clamp electrophysiology

Whole-cell patch clamp recordings were performed on hiPSC-sympathetic neurons (day 40-77). We employed an AxoPatch 200B amplifier (Molecular Devices), digitized with Digidata 1550B with a SliceScope pro upright microscope. Borosilicate glass capillaries (Harvard GC150TF-10, #30-0066) were pulled using a Narishige puller at a resistance of 4.5-5.5 MΩ and filled with (in mM): 130 K-Gluconate, 2 NaCl, 5 EGTA, 0.5 CaCl_2_, 2 MgCl_2_, 10 HEPES, 2 ATP-Na, 0.5 GTP-Na, 5 Phosphocreatine-Na (pH 7.4 with KOH; Osm 275). Cells were continuously perfused at room temperature (24-26°C) with an extracellular solution (mM): 140 NaCl, 5 KCl, 1 MgCl_2_, 2 CaCl_2_, 10 Glucose, 10 HEPES (pH 7.4 with NaOH). The liquid junction potential was calculated using the Clampex Junction Potential Calculator as −15mV and adjusted offline. All recordings were acquired and analysed using the pClamp software v11.3. Input Resistance (R_i_) was analysed in Current Clamp mode, recording the voltage response to +10mV increments from −100 to +20mV with a 500ms current injection at a 1Hz frequency. Evoked action potentials were measured upon 500ms current injection at 0.1Hz frequency, from −80 to 240pA with stepwise 20pA increment, or until 2x rheobase was reached. Resting membrane potential was calculated as the baseline average of the 200ms before the stimulus. Rheobase was defined as the minimum injected current that elicited an action potential. Cells that did not produce an action potential or with a resting membrane potential above −40mV were excluded for analyses. Neurons were classified depending on the firing pattern using the “evoked action potential” protocol similar to (Luther & Birren, 2009): “Phasic 1”, if they only fired one action potential, “Phasic 2” if they evoked 2-3 action potentials and only in the first half of the protocol, “Tonic” if they evoked multiple action potentials across the protocol.

The following drugs were applied in the recording bath using a gravity-fed perfusion system: Nicotine ditartrate (#3546/50, Bio-techne) L-(-)-Norepinephrine (+)-bitartrate salt monohydrate (#A9512, Sigma-Aldrich), UK 14,304 (10μM, #0425 Bio-techne), Carbachol (2μM, #C4382 Sigma-Aldrich), ML297 (1μM and 10μM #5380 Bio-techne) and Tertiapin Q (100nM, #1316 Bio-techne) was applied with 10μM UK 14,304 or 2μM Carbachol.

Changes in resting membrane were assessed following 1 minute of application and 2 minutes for the washout. For rheobase, input resistance and for the input/output curve, the protocol for evoked action potential was initiated following 3 minutes of drug application and 6 minutes following washout. The effect of the drugs on the input resistance was analysed using the evoked AP protocol analysing the sweeps from −80pA to +20pA. The effect of nicotine on resting membrane potential or action potential generation was assessed within 60 second following application. To test the involvement of the Gα_i/o_ subunit in the muscarinic pathway, cells were treated for 19-24hours with pertussis toxin (PTx, #3097 Bio-techne) (100ng/mL in neuronal media) and the effect of 2μM carbachol was assessed as explained above. A similar approach was employed in HEK cells expressing GIRK and muscarinic receptors (Leaney *et al*., 2001) and in peripheral neurons (Quallo *et al*., 2017).

### Ca^2+^ imaging and analyses

Coverslips of D40-45 hiPSC-sympathetic neurons were incubated in extracellular solution (ECS, composition as described for whole cell patch-clamp electrophysiology) supplemented with 2.63μM of the ratiometric calcium indicator Fura-8 AM (21055, AAT Bioquest) and 1μM probenecid (P8761, Sigma) for 2 h at 37°C and 5% CO_2_.

Following incubation, coverslips were transferred to a recording chamber and continuously superfused with ECS at a flow rate of 3ml/min. Cells were imaged with an OptoFrame microscope (Cairn Research) equipped with an OptoLED illumination system (Cairn Research), using excitation wavelengths of 365nm and 405nm, and emission wavelength > 510nm. Image acquisition was performed using MetaFluor Fluorescence Ratio Imaging Software (Molecular devices).

Ca^2+^ imaging experiments were conducted at 20.5 ± 1°C. Cells were continuously perfused with ECS or drugs diluted in ECS, nicotine (1 or 10μM, nicotine ditartrate, #3546/50, Bio-techne) or carbachol (2μM #C4382 Sigma-Aldrich). Data were normalised to cells responding to a high-potassium solution (referred as 50mM KCl throughout the text) to confirm live and excitable cells containing (in mM): 95 NaCl, 50 KCl, 10 Glucose, 10 HEPES, 2 CaCl_2_, 1 MgCl_2_, (pH = 7.4, adjusted with NaOH).

Cells responsive to a final 50mM KCl challenge were identified and selected for analysis. Regions of interest were defined manually using Metafluor, and the ratio of fluorescence emission after alternate excitation at 365 and 405 nm were exported to Excel (Microsoft). This ratio was employed as an index of evoked intracellular Ca^2+^ ([Ca^2+^]_i_) responses.. The coefficient of variation over a 3-minute baseline period was calculated for each cell. Stimulus-responsive cells were defined as those showing an increase in fluorescence ratio greater than 3 times the baseline coefficient of variation. Data were plotted and analysed in Prism 10 (GraphPad).

### Single cell RNA sequencing analyses

We used the single cell RNA sequencing neuronal cluster of the dataset of mouse thoracic sympathetic ganglia GSE78845 (Furlan *et al*., 2016). Genes were visualised as dot plot using R, the size of the dot shows the percentage of expressing cells, and the colour the average of expression (molecules per cell).

### Statistical analyses

All data were represented and analysed using Prism 10 (GraphPad). In each graph we reported the number of cells (n) and number of batches (N) that were employed. Data were assessed for normality distribution using the D’Agostino & Pearson test. For the experiments where the effect of a drug was assessed before/after application, paired tests were performed, the specific statistical test employed is described in each Figure. Statistical significance was defined as *p* ≤0.05. All data are reported and shown as mean ± SD.

## RESULTS

### hiPSC-sympathetic neurons reproduce key features of native neurons and present mature electrophysiological properties

In order to explore modulatory pathways in human sympathetic neurons, we generated hiPSC-derived sympathetic neurons (Figure 1A) using an hiPSC line from a healthy donor. We followed the protocol published by (Winbo *et al*., 2020), and, as explained in our Method section “Generation of hiPSC-sympathetic neurons”, we introduced a neuronal enrichment step (Figure 1B): AraC treatment to reduce mitotic non-neuronal cells. To verify the generation of sympathetic neurons we employed a combination of RT-qPCR and immunocytochemistry. The undifferentiated cells were tested for the expression of the pluripotency markers (*NANOG*, *POU5F1*) as a quality control, which reduced upon differentiation (Figure 1D). To confirm the successful differentiation towards peripheral sympathetic lineage we assessed the expression for the following markers: Peripherin (*PRPH*, marker for peripheral neurons), paired-like homeobox 2B (*PHOX2B*, autonomic marker), Dopamine β -hydroxylase, (*DBH*, the enzyme that catalyses the conversion of dopamine into norepinephrine, sympathetic marker) (Bos *et al*., 2025), Neuropeptide Y (*NPY*, sympathetic co-transmitter) and *CHRNA3* (nicotinic receptor subunit highly expressed in sympathetic ganglia). All these transcripts were expressed at higher levels in hiPSC-derived sympathetic neurons, compared to undifferentiated cells (Figure 1C). Immunocytochemistry confirmed that the majority of differentiated cells expressed autonomic and sympathetic markers (66% of DAPI positive cells expressed the sympathetic, TH, and 66% the autonomic, PHOX2B, markers) (Figure 1E-F).

**Figure 1.**
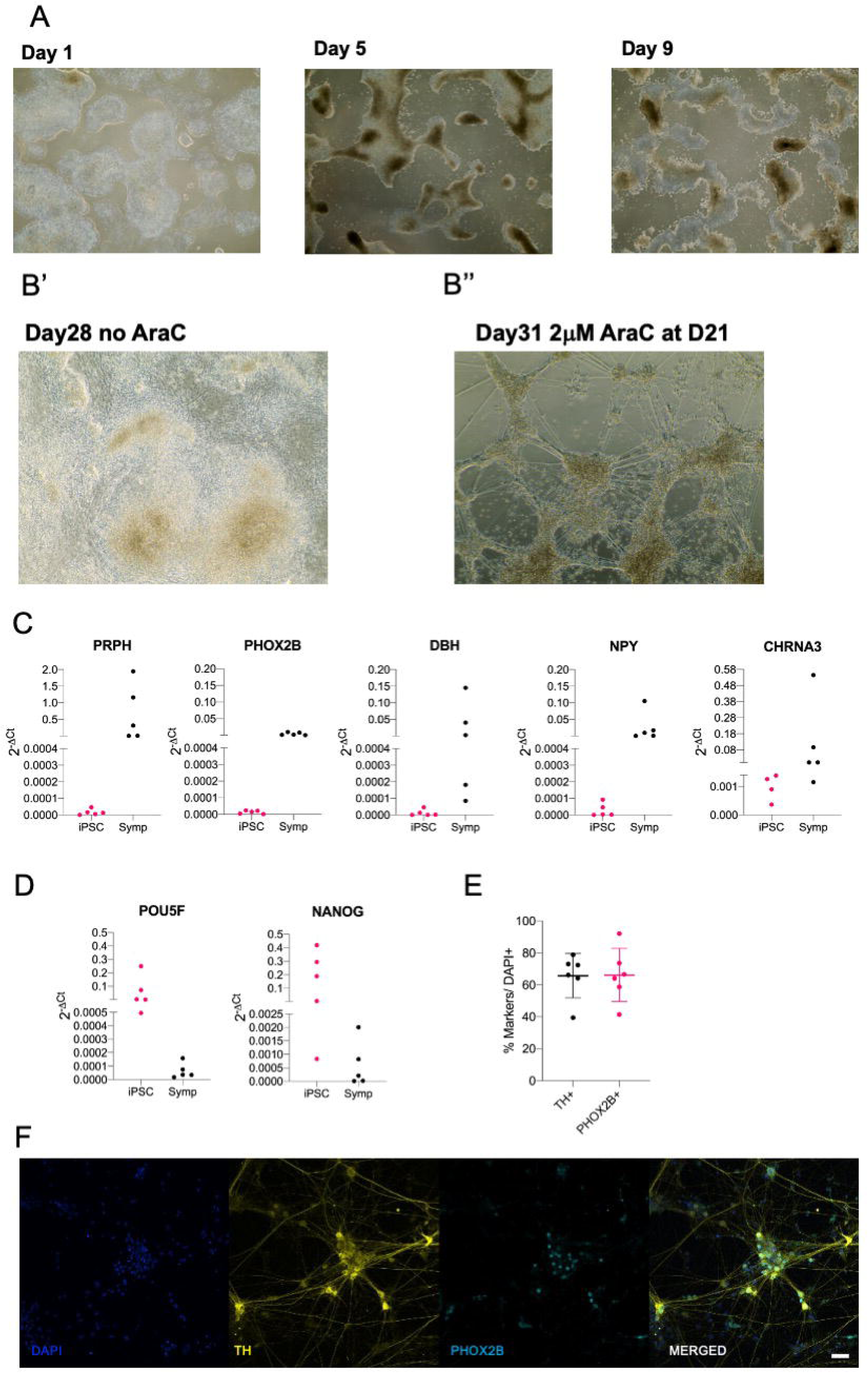
Human induced stem cells successfully differentiate into sympathetic neurons. (**A**) Phase contrast images of cells during differentiation. (**B**) Sample image of cells in the absence of AraC (**B’**) or following AraC treatment (treated at Day 21 for 72h) (**B’’**). (**C**) hiPSC-derived sympathetic neurons (Symp) express the peripheral neuron marker peripherin (*PRPH*), the autonomic marker *PHOX2B*, the sympathetic marker dopamine β -hydroxylase (*DBH*), Neuropeptide Y (*NPY*) and the ganglionic nicotinic subunit gene *CHRNA3*, compared to undifferentiated hiPSCs (iPSC). (D) Expression of the the pluripotency markers *POU5F* and *NANOG* in undifferentiated hiPSCs and in hiPSC-derived sympathetic neurons. (E) Percentage of DAPI positive cells expressing PHOX2B and TH. (F) Representative image of hiPSC-sympathetic neuron stained for PHOX2B and TH Scale bar: 50*μ*m Each dot in C and D represents a distinct differentiation batch (n=5 undifferentiated hiPSCs and hiPSC-sympathetic neurons). Each dot in E illustrates the average of two coverslip per batch, n=6.

Next, we assessed the physiological properties of the differentiated hiPSC sympathetic neurons using a combination of Ca^2+^-imaging and whole cell patch clamp electrophysiology. Sympathetic neurons are activated by acetylcholine, released from the preganglionic neurons, which activates the nicotinic receptors on the soma of sympathetic ganglia (Habecker *et al*., 2024). Given the key role of nicotinic receptors in sympathetic neurons, we applied two concentrations (1μM and 10μM) of the selective agonist nicotine (Figure 2A-C). We analysed a total of 675 cells (6 coverslips/2 batches) that responded to 50mM KCl and found that 89% of them responded to nicotine. Nicotine (1µM) evoked [Ca^2+^]_i_-responses in 68% of excitable cells, whereas 21% only responded to 10µM nicotine.

**Figure 2.**
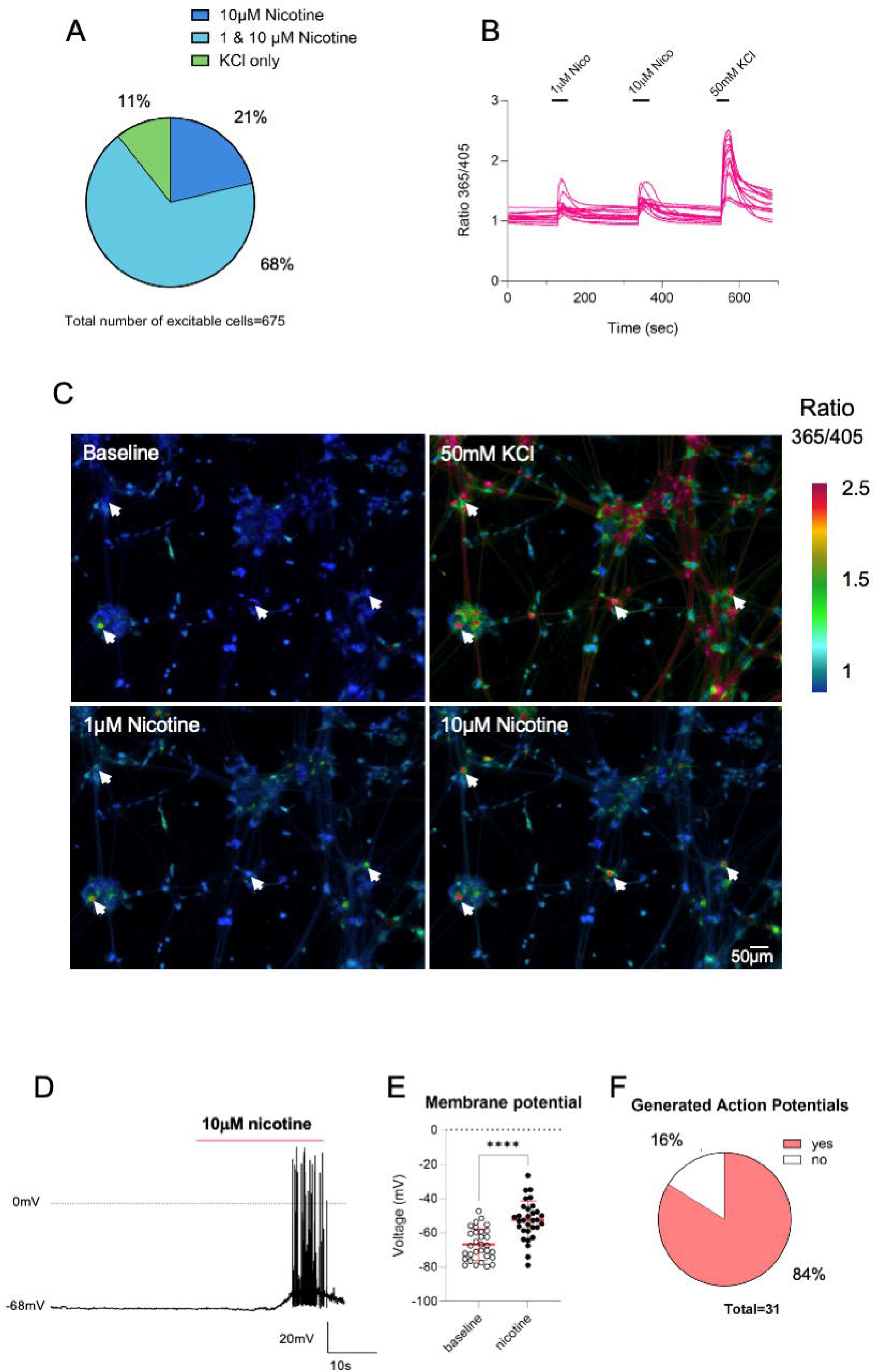
The majority of excitable cells respond to nicotine. (**A**) Distribution of excitable cells (defined by responses to 50mM KCl) responding to the application of 1*μ*M and 10*μ*M nicotine in Ca^2+^ imaging experiments (Fura-8) (n=675, 6 coverslips, N=2. (**B**) Representative example traces of changes in [Ca^2+^]_i_ in hiPSC-derived sympathetic neurons upon the application of 1*μ*M and 10*μ*M nicotine and 50mM KCl. (C) Representative pseudo-coloured images illustrating [Ca^2+^]_i_ at baseline and by stimulation with 1 *μ*M and 10 *μ*M nicotine and 50mM KCl in fura-8 loaded cells. Scale bar: 50*μ*m. Arrows indicates cells that responded to 1,10*μ*M nicotine and 50mM KCl. (D) Sample trace of whole-cell patch clamp recordings (current clamp) showing hiPSC-derived sympathetic neurons depolarization and action potential firing upon 10*μ*M Nicotine application. (**E**) Effect of nicotine on resting membrane potential (RMP), two-tailed paired t test *p*<0.0001 (baseline: −66.80±9.116mV; nicotine: −52.54±11.19mV); (**F**) Percentage of neurons that resulted in the generation of action potential upon nicotine application (n=31, N=5). Red Bar in D represents the application of nicotine

Whole cell patch clamp current clamp recordings (Figure 2D-F) demonstrated that application of 10μM nicotine elicited membrane depolarisation (Figure 2D-E) (+14.26±9.53mV) and resulted in action potential generation in 84% of cells, with the effect occurring within 20-60 seconds of drug bath application. A similar proportion to the 89% that responded with an increase [Ca^2+^]_i_ (Fig. 2A-C).

Next, we analysed the firing patterns of the hiPSC-sympathetic neurons across 6 batches (123 cells) (Figure 3). Overall, the cells exhibited tonic (80%) and phasic (Phasic 1 18% and Phasic 2 2%) firing profiles (Figure 3A, B), consistent with previously reported data on hiPSC-sympathetic neurons (Winbo *et al*., 2020; Li *et al*., 2026). Our data are also consistent with the sub-classification of rodent sympathetic ganglia further confirming that the hiPSC-sympathetic neurons mimic the native counterparts (Luther & Birren, 2009; Davis *et al*., 2020; Verkerk *et al*., 2025).

**Figure 3.**
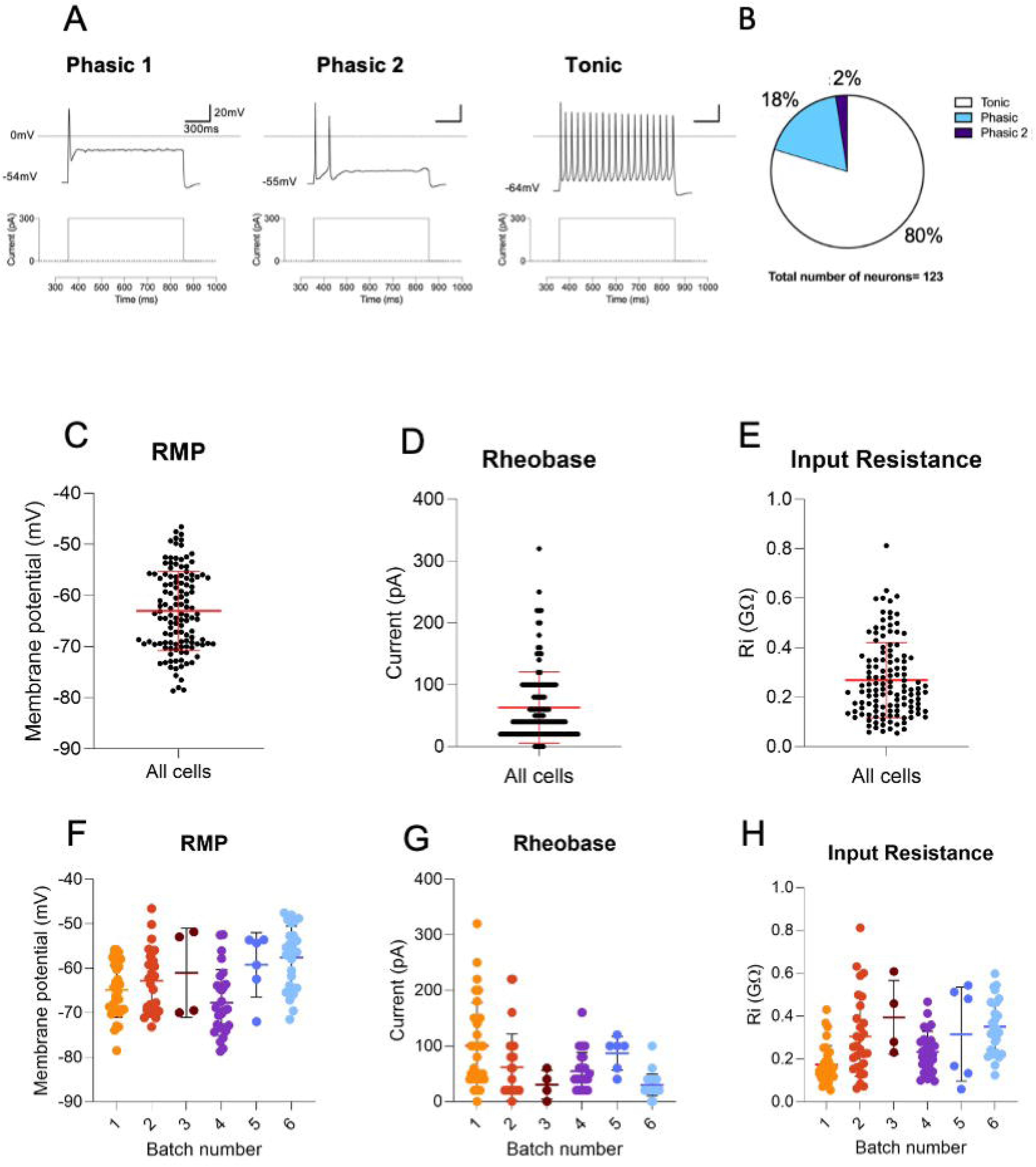
Electrophysiological characterisation of hiPSC-derived sympathetic neurons. **(A**) Sample traces of phasic (phasic 1 and 2) and tonic firing hiPSC-derived neurons upon the same current stimulation (300pA) and (**B**) percentage of tonic (80%) and phasic (Phasic 1 18% and Phasic 2 2%) neurons (n=123/N=6). Summary of (**C**) Resting membrane potential (RMP: −63.03±7.77mV), (**D**) Rheobase (63.33±57.97pA), and (**E**) Input Resistance (0.269±0.152GΩ) and analysed per batch (**F-H**) (n=123/N=6). Batch 1 day 44-62, Batch 2 day 42-77, Batch 3 day 47-52, Batch 4 42-56, Batch 5 day 48-50, Batch 6: day 41-60.

Differentiated cells showed resting membrane potential (−63.03±7.77mV), input resistance (0.269±0.152GΩ) and rheobase (63.33±57.97pA) (Figure 3C-H) in line with previously published work on hiPSC-sympathetic neurons (Winbo *et al*., 2020) and rat stellate ganglia (Davis *et al*., 2020). Overall, our data therefore demonstrate that hiPSC-sympathetic neurons from day 40 express key autonomic and sympathetic markers, and express functional nicotinic receptors and physiological properties characteristic of their native counterparts.

### Adrenergic receptors modulate the activity of hiPSC-sympathetic neurons

Adrenergic receptors are known to modulate sympathetic neurons, but the mechanisms that regulate neuronal excitability have not been fully elucidated. Firstly, we analysed the neuronal cluster of the single cell RNAseq (GSE78845) (Furlan *et al*., 2016) dataset of mouse sympathetic thoracic ganglia to investigate the expression profile of adrenergic receptors in sympathetic neurons. Our analyses showed high expression of the α2 adrenergic receptor genes (*Adra2a*, *Adra2b* and *Adra2c*), some expression of the β 2 adrenergic receptor gene but essentially no expression of α1 adrenergic receptor genes (*Adra1a*, *Adra1b*, *Adra1c*, *Adra1d*) (Figure 4A).

**Figure 4.**
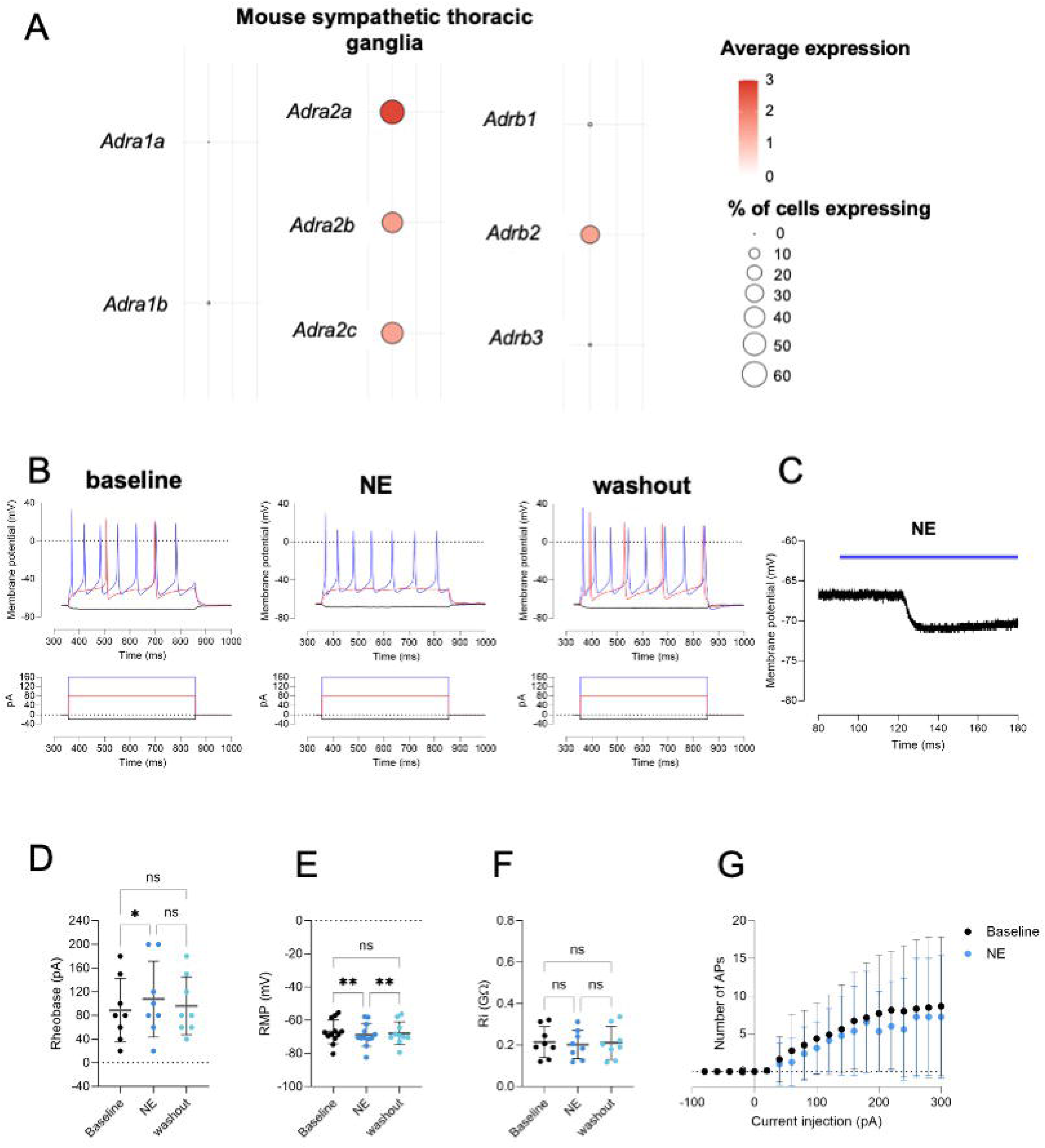
Norepinephrine (100μM) reduces the excitability of hiPSC-derived sympathetic neurons. >**A**) Expression of adrenergic receptor genes in mouse thoracic sympathetic ganglia (scRNAseq dataset GSE78845). Colour scale shows the average of expression (molecules per cells), size of the dot percentage of cells. (**B**) Sample trace of evoked action potential at baseline, upon norepinephrine (100*μ*M) application and washout. (**C**) Sample trace of the effect of norepinephrine (100*μ*M) on resting membrane potential. Graphs summarising the effect of norepinephrine on (**D**) Rheobase (baseline: 88.75±53.84pA, NE: 107.5±64.09pA; 96.25±48.97pA: Repeated Measures One-way ANOVA *p*=0.037; Tukey’s multiple comparisons test: baseline vs NE *p*=0.0266, baseline vs washout *p*=0.3796, NE vs washout *p*=0.34, n=8/N=2). (**E**) Resting membrane potential (baseline: −66.99±7.188mV, NE: −68.76±6.918mV; washout: −67.92±6.885mV; mixed effects model *p*=0.0011, Tukey’s multiple comparisons tests: baseline vs NE *p*=0.0016, baseline vs washout *p*=0.7991, NE vs washout *p*=0.0085 n=12/N=2 for baseline and NE, n=10/N=2 for washout) (**F**) Input Resistance (baseline: 0.215±0.075GΩ, NE: 0.203±0.069GΩ, washout: 0.211±0.080GΩ; Repeated Measures One-way ANOVA *p*=0.2685, n=8/N=2 (**G**) Frequency of Action potential induced by an increasing stepwise current injection. Mixed effect analysis: Number of Action potentials upon current injection *p*=0.049, upon drug application *p*=0.795, current injection x drug application *p*=0.334, n=8/N=2.

Using whole-cell patch clamp recordings, we examined the effects of the sympathetic neurotransmitter and adrenergic receptor agonist, norepinephrine on hiPSC-sympathetic neurons. We analysed the drug effects on electrophysiological parameters involved in neuronal excitability: resting membrane potential, rheobase, firing frequency upon current injection and input resistance. Norepinephrine (100μM, NE) reduced the excitability of hiPSC-derived sympathetic neurons, evident by the increase in rheobase (Repeated Measures one-way ANOVA *p*=0.0370, Tukey’s multiple comparisons tests: baseline vs NE *p*=0.0266, baseline vs washout *p*=0.3796, NE vs washout *p*=0.34) (Figure 4B, D), hyperpolarisation (mixed effects model *p*=0.0011, Tukey’s multiple comparisons tests: baseline vs NE *p*=0.0016, baseline vs washout *p*=0.7991, NE vs washout *p*=0.0085) (Figure 3C, E) with both effects fully reversed upon washout. However, norepinephrine application did not affect the input resistance significantly (Repeated Measures one-way ANOVA *p*=0.2685) (Figure 4F) nor firing frequency (Figure 4G) evoked by stepwise current injection. Given that α2 adrenergic (α2ARs) receptors are the highest expressed subtypes in mouse sympathetic ganglia, and that α2AR predominantly signal via Gα_i_, shown to modulate sympathetic activity (Lipscombe *et al*., 1989; Trendelenburg *et al*., 2001) and neuronal firing (Nimitvilai *et al*., 2017) we hypothesised that the effect of NE could be mediated by α2ARs. We first evaluated the expression levels of the α2ARs genes in our human system (Figure 5A). Consistent with the mouse transcriptomics data, hiPSC-derived sympathetic neurons expressed α2ARs with *ADRA2A* higher than either *ADRA2B* or *ADRA2C*. We employed the α2ARs selective agonist UK14304 (10μM) and found that it elicited similar responses to norepinephrine. UK14304 increased the rheobase (Mixed effects models *p*=0.0448; Tukey’s multiple comparisons tests: baseline vs UK14304 *p*=0.0219, baseline vs washout p=0.7324, UK14304 vs washout *p*=0.0166) (Figure 4B, D), produced membrane hyperpolarization (Mixed effects models *p*=0.0125; Tukey’s multiple comparisons tests: baseline vs UK14034 *p*=0.0065, baseline vs washout *p*=0.7604, UK14304 vs washout *p*=0.0521; Figure 5C and E) and reduced input resistance (Mixed effects models *p*=0.030, Tukey’s multiple comparisons tests: baseline vs UK14304 *p*=0.0321, baseline vs washout *p*=0.4955, UK14304 vs washout *p*=0.0951) (Figure 5F). Moreover, UK14304 also had a significant effect on firing frequency upon stepwise current injection (mixed effects analysis: current injected *p*<0.0001, Treatment *p*=0.0137, current injected x Treatment *p*=0.0008 and a significant reduction of firing frequency when 140pA was injected *p*=0.0020). Overall, our data show that norepinephrine reduces the excitability of the neurons with this effect mimicked by the α2 adrenergic receptors agonist.

**Figure 5.**
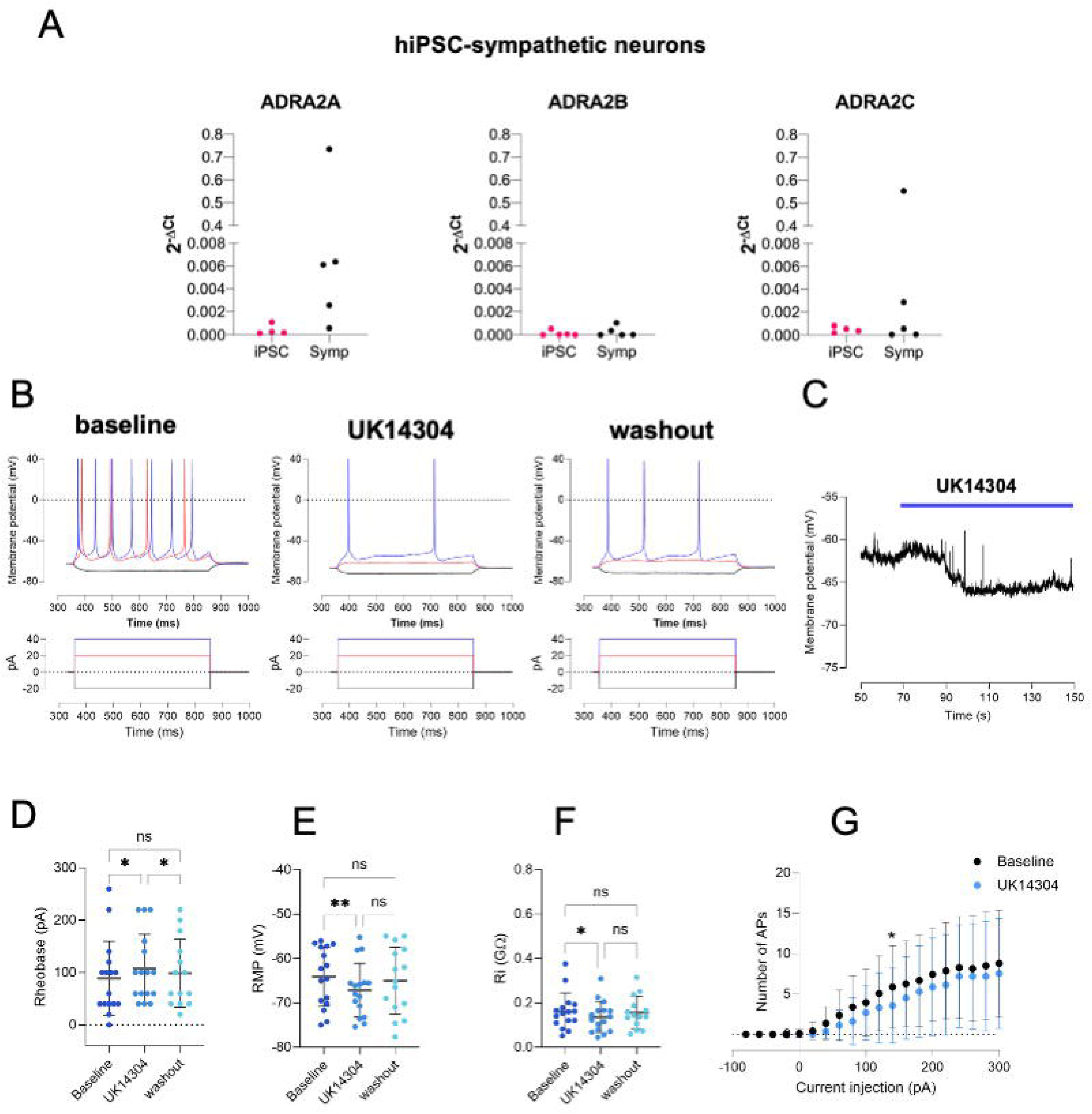
The α2 adrenergic receptor agonist UK14304 (10μM) reduces the excitability of hiPSC-sympathetic neurons. (**A**) RT-qPCR of α2 adrenergic receptors transcripts in hiPSC-sympathetic neurons and undifferentiated hiPSCs. Each dot in C and D represents a distinct differentiation batch (n=5 undifferentiated hiPSCs and hiPSC-sympathetic neurons. (**B**) Sample trace of evoked action potential at baseline, in the presence of 10*μ*M UK14304 and after washout. (**C**) Sample trace of the effect of 10*μ*M UK14304 on resting membrane potential. Graphs summarising the effect of UK14304 on (**D**) rheobase (baseline: 88.75±71.17pA; UK14304 107.15±65.68pA; washout, 98.57±64.91pA; Mixed effects models *p*=0.0448; Tukey’s multiple comparisons tests: baseline vs UK14304 *p*=0.0219, baseline vs washout *p*=0.7324, UK14304 vs washout *p*=0.0166. (**E**) Resting membrane potential (baseline:-64.12±6.672mV, UK14304 –67.07±6.027mV, washout:-64.99±7.571mV; Mixed effects models *p*=0.0125; Tukey’s multiple comparisons tests: baseline vs UK14034 *p*=0.0065, baseline vs washout *p*=0.7604, UK14304 vs washout *p*=0.0521) (**F**) Input resistance (baseline 0.162±0.082GΩ UK14304: 0.136±0.070GΩ washout:0.157±0.072GΩ; Mixed effects models *p*=0.030, Tukey’s multiple comparisons tests: baseline vs UK14304 *p*=0.0321, baseline vs washout *p*=0.4955, UK14304 vs washout *p*=0.0951. (**G**) Frequency of action potential induced by an increasing stepwise current injection (Mixed effect analysis: Number of Action potentials upon current injection *p*<0.0001, upon drug application *p*=0.0137, current injection x drug application *p*=0.0008; with a significant reduction of firing frequency when 140pA was injected *p*=0.0020. (n=16, N=2 for baseline and UK14304, n=14, N=2 for washout).

### Muscarinic receptors inhibit the activity of hiPSC-sympathetic neurons

Muscarinic receptors modulate the activity of sympathetic neurons (Trendelenburg *et al*., 2003; Jani *et al*., 2025). To identify which receptor subtypes are most likely to be responsible, we first assessed the expression levels of muscarinic receptor genes in the mouse sympathetic ganglia transcriptomic dataset (GSE78845) (Furlan *et al*., 2016). As shown in Figure 6A the highest expressed gene was *Chrm2* followed by *Chrm1*, with *Chrm3*, *Chrm4* and *Chrm5* virtually not expressed. The qPCR analyses confirmed that *CHRM2* was expressed at higher levels than *CHRM1* also in hiPSC-sympathetic neurons (Figure 6B). Muscarinic receptors in sympathetic ganglia have been reported to have an excitatory effect through M1 (Shapiro *et al*., 2001) or inhibitory through M2 and M4 (Shapiro *et al*., 2001), M2 in cardiac innervating neurons (Trendelenburg *et al*., 2003; Jani *et al*., 2025) and some reports suggesting an inhibitory role also through the Gα_q_- coupled M1 (Delmas *et al*., 1998). We first assessed whether the muscarinic agonist carbachol (2μM) activated hiPSC-derived sympathetic neurons employing intracellular [Ca^2+^]_i_ as an index of neuronal activation (Fig 6C). Carbachol application was followed by nicotine (10μM) to validate the sympathetic identity of the cells recorded. Using a total of 813 cells (6 coverslips/2 batches), we found that carbachol did not elicit increase in [Ca^2+^ ]_i_ in 98% of 50mM KCl responders. In this set of coverslips, nicotine (10μM) elicited increase in [Ca^2+^]_i_ in 96% of KCl-responsive cells. Given that carbachol did not elicit an overall activation of the neurons, we employed whole cell patch clamp electrophysiology (current clamp) as a more sensitive approach to detect any other forms of modulatory activity. Carbachol application resulted in an overall inhibition of neuronal activity similar to norepinephrine (Figure 4) and UK14304 (Figure 5). Overall, it increased the rheobase (*p*=0.0177; Tukey’s multiple comparisons tests baseline vs CCH *p*=0.0004, baseline vs washout *p*=0.5881, CCH vs washout *p*=0.0029) (Figure 7C, E), hyperpolarised the neurons (*p*=0.0011; Tukey’s multiple comparisons tests baseline vs CCH *p*=0.0063, baseline vs washout *p*=0.9739, CCH vs washout *p*=0.0222) (Figure 7D and F) but did not have a significant effect on input resistance (*p*=0.0526) nor on firing frequency upon stepwise current injection using a mixed effect models. Of note, in two cells with a higher rate of spontaneous action potentials, carbachol application reduced their spontaneous activity, and this effect was reversed after washout (sample trace shown in Figure 7G and H).

**Figure 6.**
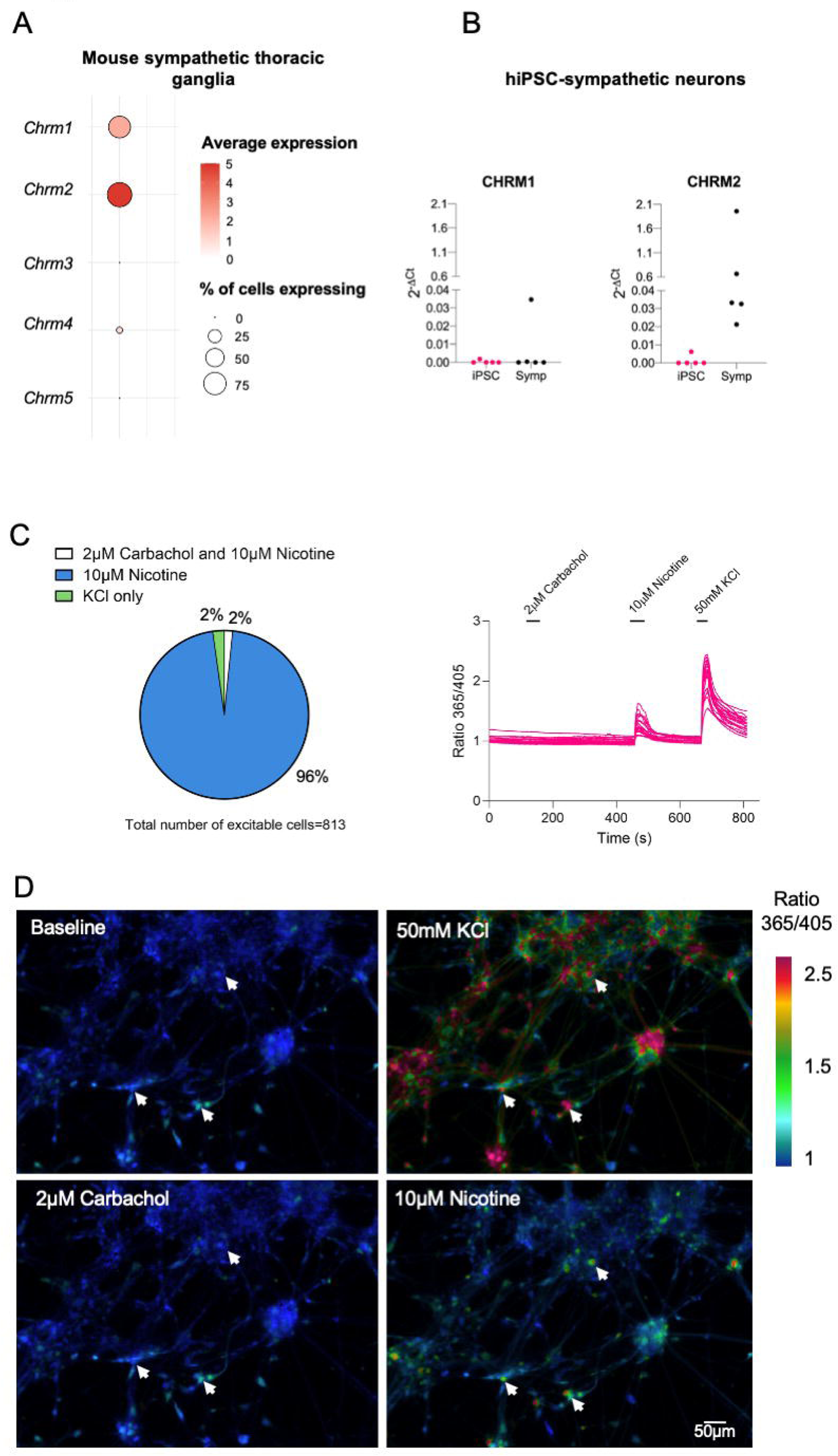
The muscarinic agonist Carbachol (2μM) does not increase [Ca^2+^]_i_ in hiPSC-sympathetic neurons. (**A)** Expression of muscarinic receptor genes in mouse thoracic sympathetic ganglia (scRNAseq dataset GSE78845). (**B**) RT-qPCR for *CHRM1* and *CHRM2* of hiPSC sympathetic neurons compared to undifferentiated hiPSCs. Each dot represents a distinct differentiation batch (N=5 undifferentiated hiPSCs and hiPSC-sympathetic neurons). (**C**) The left-hand panel shows the proportions of excitable cells (responsive to 50 mM KCl) that resulted in increase in [Ca^2+^]_i_ upon 2*μ*M carbachol application, followed by 10*μ*M nicotine (as a positive control, 96% of excitable cells), and 50mM KCl. The right-hand panel shows representative traces of [Ca^2+^]_i_ responses in hiPSCs-derived sympathetic neurons upon carbachol, nicotine (10*μ*M) and KCl (50mM) application. Carbachol (2*μ*M) application did not elicit a Ca^2+^ response in 98% of cells (activated by 50mM KCl) while nicotine (10*μ*M) elicited [Ca^2+^]_i_ increase in 96% of KCl-responders; n=813, 6 coverslips, N=2. (**D**) Representative pseudo-coloured images illustrating [Ca^2+^]_i_ at baseline, following 2*μ*M Carbachol, 10*μ*M nicotine and 50mM KCl in fura-8 loaded cells. Arrows indicate cells that responded to 10*μ*M nicotine and 50mM KCl. Scale bar = 50*μ*m.

**Figure 7.**
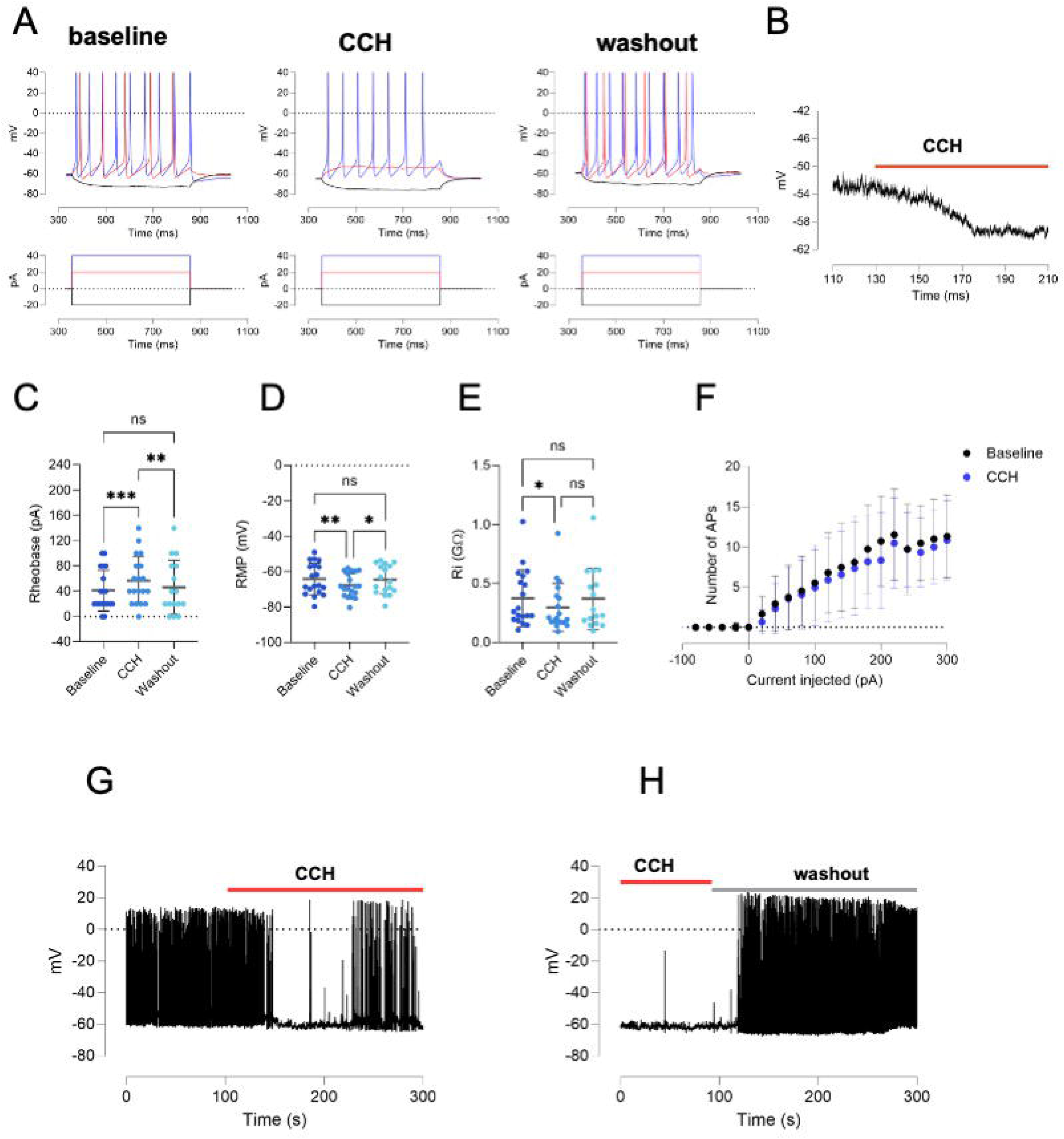
The muscarinic agonist Carbachol (2μM) reduces the excitability of hiPSC-sympathetic neurons. (**A**) Sample trace of evoked action potential at baseline, in presence of 2*μ*M Carbachol and upon washout. (**B**) Sample trace of the effect of Carbachol on resting membrane potential. Graphs summarising the effect of carbachol on (**C)** rheobase (baseline: 41.05±32.30pA, CCH: 56.84±37.87pA, washout: 46.25±42.41pA; RM Mixed effect model *p*=0.0177; Tukey’s multiple comparisons tests baseline vs CCH *p*=0.0004, baseline vs washout *p*=0.5881, CCH vs washout *p*=0.0029) (**D)** Resting membrane potential (baseline:-64.09±8.868mV, CCH:-67.68±6.621mV; washout:-64.50±8.229mV; RM mixed model models *p*=0.0011; Tukey’s multiple comparisons tests baseline vs CCH *p*=0.0063, baseline vs washout *p*=0.9739, CCH vs washout *p*=0.0222) (**E**) Input resistance (baseline:0.375±0.239 GΩ, CCH:0.298±0.203GΩ, washout: 0.371±0.258GΩ; RM mixed effect models: *p*= *p*=0.0526) (**F**) Frequency of action potential induced by an increasing stepwise current injection (Mixed effect analysis: Number of Action potentials upon current injection *p*<0.0001, upon drug application *p*=0.7091, current injection x drug application *p*=0.8716) (n=19, N=3 for baseline and CCH; n=16, N=3 for washout). Sample trace of (**G**) carbachol inhibition of spontaneous action potential, reversed upon washout (**H**).

The muscarinic receptors M1 and M2 typically couple to the Gα_q/11_ and Gα_i/o_ subfamilies respectively (Shapiro *et al*., 2001). In order to identify which Gα-pathway is engaged by CCH in hiPSC-sympathetic neurons, we employed pertussis toxin (PTx), which catalyses the ADP ribosylation of the Gα_i/o_ subunits thereby abrogating the interaction between Gα_i/o_ and their cognate G protein coupled receptors (GPCRs) (Quallo *et al*., 2017). As shown in Figure 8, carbachol (2μM) following PTx pre-treatment (19-24h) did not elicit any discernible influence on neuronal excitability. This is evident by the lack of changes in rheobase (*p*=0.626) (Figure 8A and 8C), resting membrane potential (*p*=0.122) (Figure 8B and 8D) and input resistance (*p*=0.3343) (Figure 8E) using a mixed effect models.

**Figure 8.**
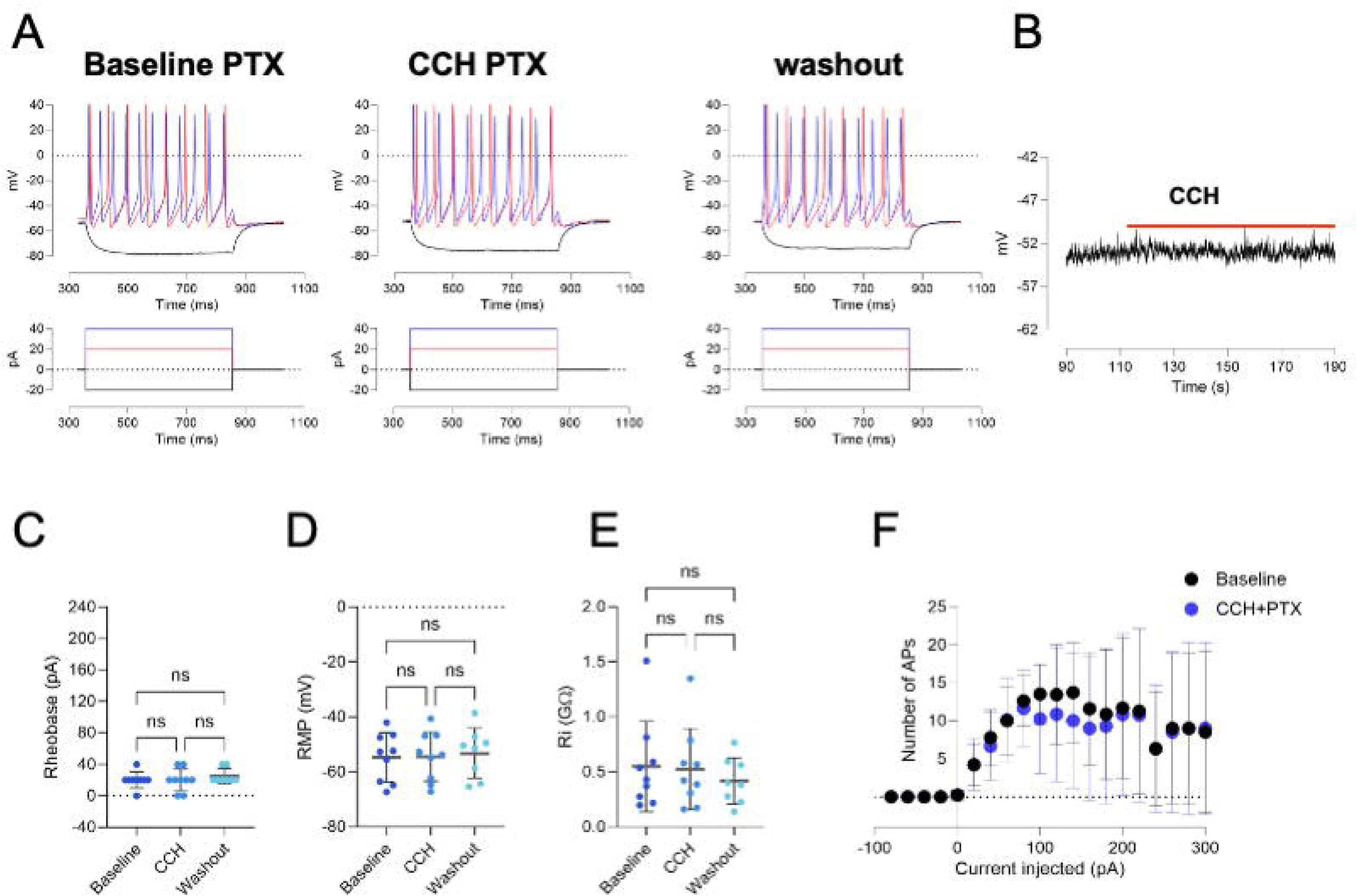
Pertussis Toxin pre-treatment inhibits Carbachol modulation. (**A**) Sample trace of evoked action potential from PTx-pretreated cells, at baseline, in presence of 2*μ*M Carbachol and upon washout. (**B**) Sample trace of the effect Carbachol application on resting membrane potential of PTx-pretreated cell. Graphs summarising the effect of carbachol on (**C)** rheobase (baseline: 20.00±10.00pA, CCH: 20.00± 14.14, washout:25.00±9.258pA; RM Mixed effects analysis *p*=0.6259) (**D)** Resting membrane potential (baseline:-54.72±8.853mV, CCH: −54.44±8.916mV, washout:-53.21±9.045mV; RM Mixed effect analysis *p*=0.122); (**E**) Input resistance (baseline: 0.549±0.4100GΩ, CCH: 0.5258±0.368GΩ, washout:0.419±0.2095GΩ RM Mixed effect analysis: *p*=0.3343); (**F**) Frequency of action potential induced by increasing stepwise current injections. Mixed effect analysis: Number of Action potentials upon current injection *p*<0.0001, upon drug application *p*=0.7561, current injection x drug application *p*=0.0073 (baseline and CCH n=9, N=1, washout: n=8, N=1)

Our data therefore show that carbachol inhibits hiPSC-sympathetic neuronal activity through a pertussis toxin-sensitive pathway, which agrees well with the known functional coupling of M2 with Gα_i/o_ and the high expression of the *CHRM2* in hIPSC-derived sympathetic neurons, as well as in native, primary sympathetic neurons.

### G-protein-coupled inwardly rectifying K^+^ channels (GIRK) are downstream effectors of adrenergic and muscarinic receptors.

Application of either α2-adrenergic or muscarinic agonists resulted in both membrane hyperpolarisation and an increased rheobase in hiPSC-sympathetic neurons (Figure 5 and 7). In other cell types, both types of G-protein coupled receptors have been reported to elicit the activation of endogenous inwardly rectifying K^+^ channels (GIRK) (Ang *et al*., 2012; Luo *et al*., 2022). Notably, direct activation of GIRK in sensory neurons produced hyperpolarisation and an increased rheobase (Brizuela *et al*., 2025). We therefore assessed whether the neuronal inhibition produced by the two ligands converged on GIRK, by using the selective blocker Tertiapin Q (100nM, TPQ) applied with UK14304 (10μM) or carbachol (2μM) (Figure 9).

**Figure 9.**
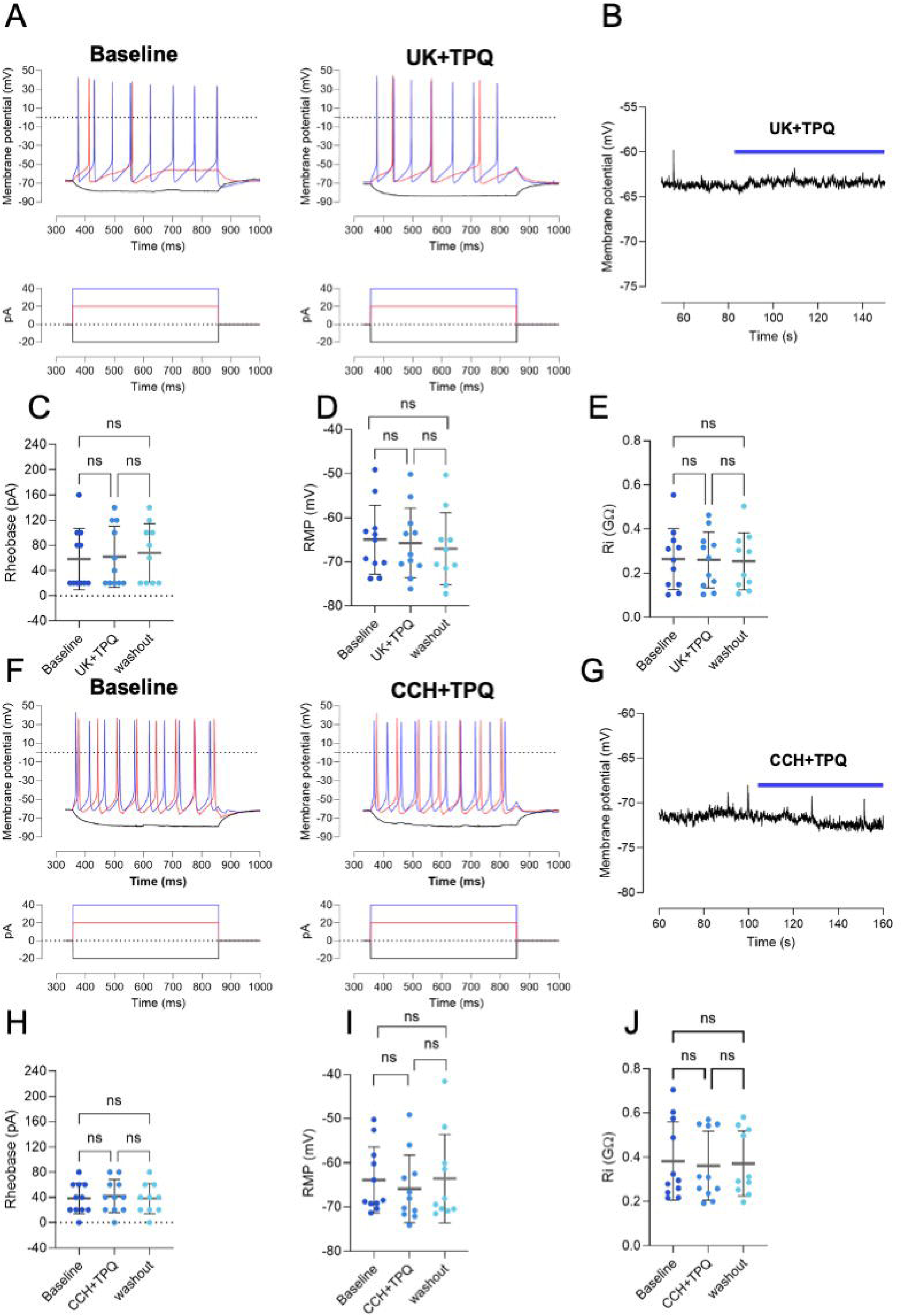
The GIRK channel blocker Tertiapin Q (100nM) abolishes the effects of carbachol and UK14304. Sample trace of evoked action potentials at (**A**) baseline and 10*μ*M UK14304 together with 100nM Tertiapin Q. (**B**) Sample trace of effect of Tertiapin Q(100nM) on 10*μ*M UK14304 on resting membrane potential. Graphs summarising the effect of application of 100nM Tertiapin Q and UK14304 on (**C)** rheobase (baseline: 58.18±48.54pA, UK+TPQ:61.82±48.54pA, washout:68.00±46.38pA; RM Mixed effect analysis: *p*=0.4050) (**D)** Resting membrane potential (Baseline: −64.96±7.819mV, UK+TPQ:-65.72±7.907mV, −66.99±8.180mV; RM Mixed effect analysis: *p*=0.0907(**E**) Input resistance (Baseline: 0.264±0.138GΩ, UK+TPQ: 0.260±0.126GΩ, washout 0.2543±0.129 GΩ; RM Mixed effect analysis *p*=0.2479. *(Baseline and UK+TPQ n=11, N=2 washout n=10, N=2)* Sample trace of evoked action potentials at (**F)** baseline and 2*μ*M Carbachol with 100nM Tertiapin Q. (G) Graphs summarising the effect of 100nM Tertiapin Q on Carbachol (2*μ*M) on (**H)** rheobase (Baseline: 38.18±24.42pA, CCH+TPQ:41.82±26.01pA, washout:38.00±23.94; RM Mixed effect analysis *p*=0.3349 (**I)** Resting membrane potential (Baseline: −63.88±7.491mV, CCH+TPQ: −65.89±7.654mV, washout: - 63.57±10.03mV; RM Mixed effect analysis *p*=0.0609, (**J**) lnput resistance baseline 0.382±0.177GΩ, CCH+TPQ: 0.362±0.156GΩ, washout 0.372±0.147GΩ; RM Mixed effect analysis *p*=0.2766 (Baseline and CCH+TPQ n=11; N=3, washout n=10, N=3).

In the presence of Tertiapin Q, the effect of UK14304 (Figure 9A-E) was abolished, and we observed no significant changes in rheobase (*p*=0.41), resting membrane potential (*p*=0907) and input resistance (*p*=0.25). Similarly, application of Tertiapin-Q reduced the inhibitory effect of carbachol (Figure 9F-J), which was evident by the lack of a significant effect on the rheobase (*p*=0.33); resting membrane potential (*p*=0.061) and input resistance (*p*=0.277). Together, these results strongly suggest that stimulation of M2 and α2 adrenergic receptors reduced the excitability of hiPSC-sympathetic neurons by activating GIRK.

### The GIRK channel opener ML297 mimics the effects of α2ARs and muscarinic agonists

To evaluate the involvement of GIRK channels in the regulation of neuronal excitability in more detail, we first assessed the expression levels of the different GIRK channel genes (*Kcnj3*, *Kcnj5*, *Kcnj6*, *Kcnj9*) in thoracic sympathetic ganglia (GSE78845). The GIRK genes detected in the datasets were *Kcnj3* and *Kcnj9* encoding respectively for GIRK1 (Kir3.1) and GIRK3 (Kir3.3) (Figure 10A). We used RT-qPCR of the hiPSC-sympathetic neurons to validate the reported expression pattern (Figure 10B) and further validated protein expression with immunocytochemistry for GIRK1 (Figure 10C).

**Figure 10.**
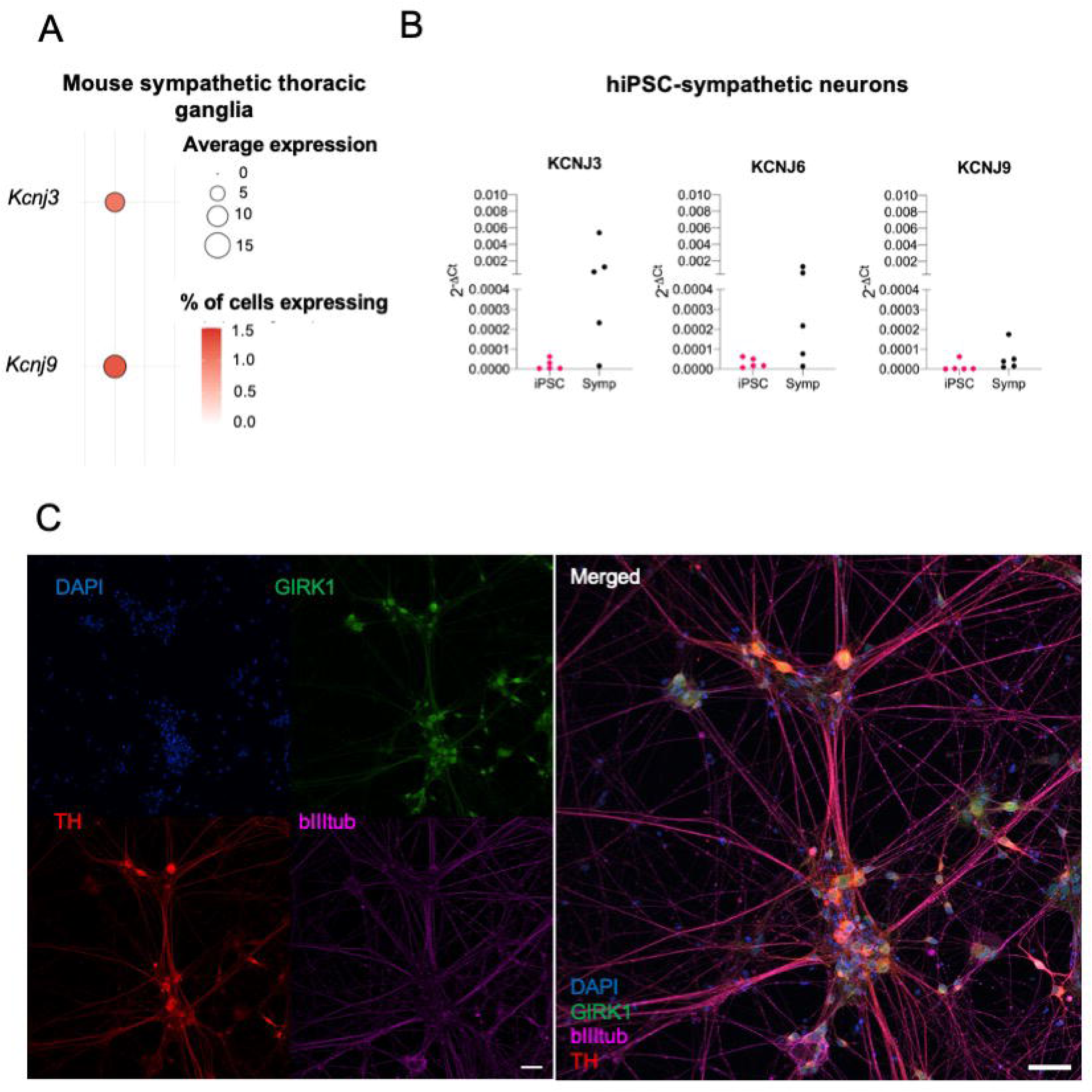
GIRK expression in sympathetic neurons. (**A**) Expression of GIRK subunits genes in mouse thoracic sympathetic ganglia (scRNAseq dataset GSE78845) only *Kcnj3* and *Kcnj9* were included in the dataset. (B) RT-qPCR of *KCNJ3*, *KCNJ6* and *KCNJ9* in hiPSC-sympathetic neurons and undifferentiated hiPSC. (**C)** Representative image of hiPSC-sympathetic neurons stained for GIRK1, TH, and beta3 tubulin (scale bar 50*μ*m)

We then employed the selective GIRK channel activator ML297 at two concentrations (1μM and 10μM), the lower of which is considered selective for GIRK1/GIRK2, while the higher would also activates GIRK3 and GIRK4 containing channels, but not GIRK2/3 complexes (Kaufmann *et al*., 2013). Application of 10μM ML297 (Figure 11A-F) resulted in a significant membrane hyperpolarisation (one-way repeated measure ANOVA *p*=0.0003; Tukey’s multiple comparisons tests baseline vs 10μM ML297 *p*=0.017, baseline vs washout *p*=0.0019, 10μM ML297 vs washout *p*=0.027), accompanied by an increased rheobase (Friedman test *p*=0.0002; Tukey’s multiple comparisons tests baseline vs 10μM ML297 *p*=0.0016, baseline vs washout *p*=0.249, 10μM ML297 vs washout *p*=0.249), a reduced input resistance (Friedman test *p*=0.0098, Tukey’s multiple comparisons tests baseline vs 10μM ML297 *p*=0.019, baseline vs washout *p*=0.041, 10μM ML297 vs washout *p*>0.99) and a significantly reduced firing frequency upon stepwise current injection (mixed effects model, current injected *p*<0.0001, Treatment *p*<0.0001, current injected x Treatment *p*=0.011 and a significant reduction of firing frequency at current 120pA *p*=0.022, 140pA *p*=0.025, 160pA *p*=0.25 and 180pA *p*=0.040). The effect of 10μM ML297 was not fully reversed upon washout. By contrast, 1μM ML297 (Figure 11 G-I) did not significantly affect neuronal excitability, with only a small reduction in the membrane potential (one-way repeated measure ANOVA *p*=0.063) and no overall effect on rheobase (Friedman test *p*=0.253) or input resistance (one-way repeated measure ANOVA *p*=0.12)

**Figure 11.**
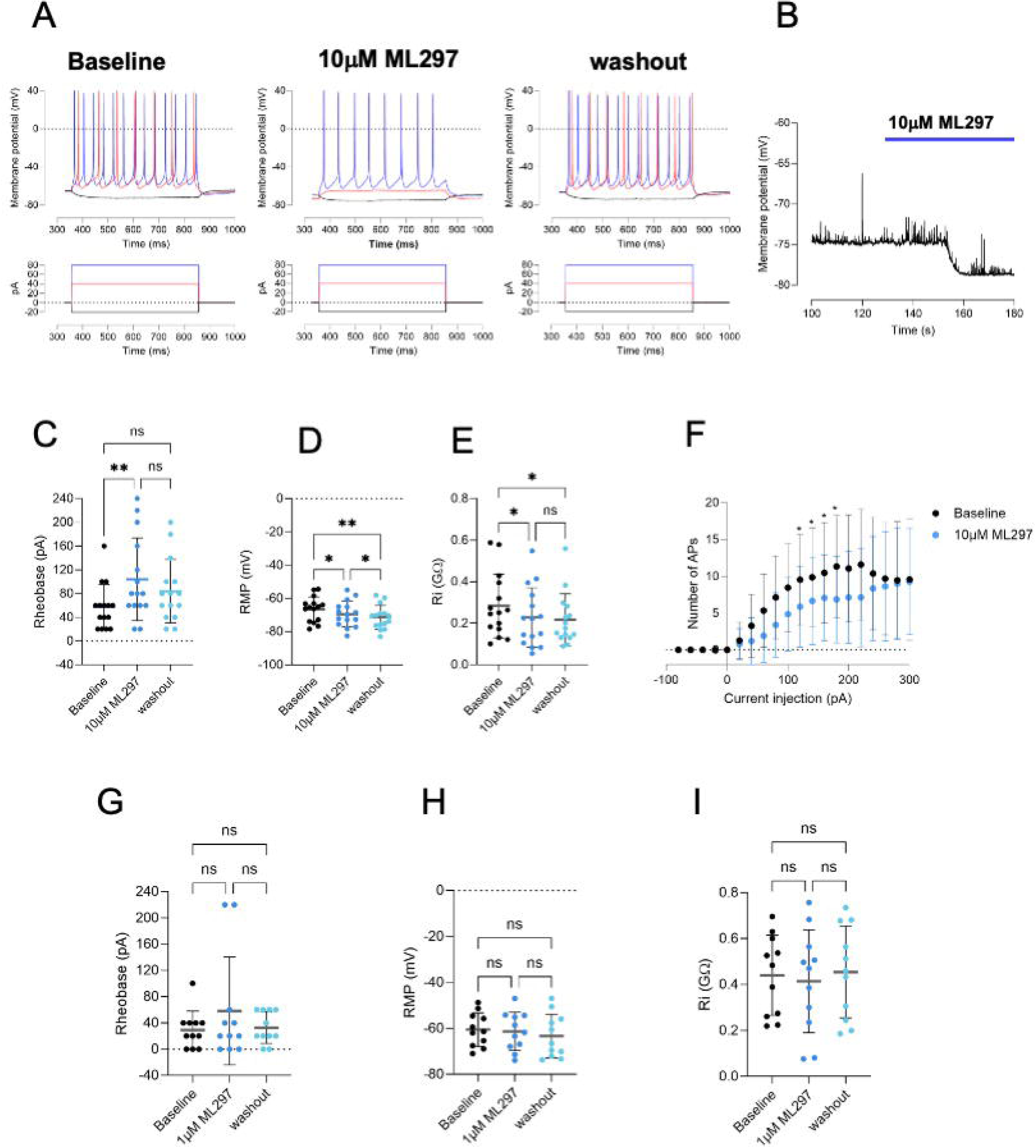
Direct activation of GIRK reduces the excitability of hiPSC-sympathetic neurons. (**A**) Sample trace of evoked action potential at baseline, in presence of 10*μ*M ML297 and upon washout. (**B**) Sample trace of the effect of 10*μ* ML297 on resting membrane potential. Graphs summarising the effect of 10*μ*M ML297 on (**C)** rheobase (baseline 57.33±38.45pA, 10*μ*M ML297 104.0±69.78pA, washout 84.00±53.56, Friedman test *p*=0.0002, Dunn’s multiple comparison test baseline vs 10*μ*M ML297 *p*=0.0016, baseline vs washout *p*=0.2485, 10*μ*M ML297 vs washout *p*=0.2485 (**D**) Resting membrane potential (baseline: −66.51±7.509mV, 10*μ*M ML297-69.42±7.733mV, washout-71.41±7.178mV; RM one-way ANOVA p=0.0003, Tukey’s multiple comparisons test baseline vs 10*μ*M ML297 *p*=0.0166, baseline vs washout *p*=0.0019, 10*μ*M ML297 vs washout *p*=0.0269 (**E**) Input resistance (baseline: 0.283±0.154GΩ, 10*μ*M ML297: 0.227±0.143GΩ, washout: 0.217±0.125GΩ (**F)** Frequency of action potential induced by an increasing stepwise current injection Number of Action potentials upon current injection *p*<0.0001, upon drug application *p*<0.0001, current injection x drug application *p*=0.0106 Graphs summarising the effects of 1*μ*M ML297 on (**G)** rheobase (baseline: 29.09±28.79pA, 1 *μ*M ML297 58.18±82.20pA, washout: 32.73±24.12pA, Friedman test p=0.2531 (**H**) Resting membrane potential (baseline: −60.53±7.317mV, 1 *μ*M ML297: −61.32±8.294, washout:-63.35±9.508mV RM one-way ANOVA p=0.0627 (**I**) Input resistance (baseline 0.440±0.174GΩ, 1 *μ*M ML297 0.414±0.224GΩ washout:0.455±0.200GΩ, RM one-way ANOVA *p*=0.1212) (10*μ*M ML297 n=15, N=3; 1*μ*M ML297 n=11, N=2).

Our data demonstrate that the direct activation of GIRK reduces the excitability of hiPSC derived sympathetic neurons. RT-qPCR on hiPSC-sympathetic neurons combined with the electrophysiological data suggest a heterogeneous GIRK channel subunits composition with a possible contribution of the GIRK3-containing channels, suggested by the reduced effect using the low concentration of 1μM ML297.

## DISCUSSION

Here, we have investigated the pathways that modulate the excitability of sympathetic neurons derived from hiPSC. We have validated the identity of the neurons as *sympathetic* based on their expression of the sympathetic markers’ tyrosine hydroxylase and dopamine β -hydroxylase, the autonomic marker *PHOX2B* and their responsiveness to nicotine. To generate hiPSC-derived sympathetic neurons, we adapted a previously published protocol and added a neuronal enrichment step using cytosine arabinoside selection. This step was based on our experience with other types of peripheral neurons (Li *et al*., 2025*b*), a similar approach was employed by other groups using other mitotic inhibitors using a different protocol for hiPSC-sympathetic neurons (Wu *et al*., 2024). As shown in Figure 1C and 1C’, without the enrichment step the non-neuronal cells dominated the culture, and we therefore suggest that the protocol we employed (Winbo *et al*., 2020) might require this step, at least for some hiPSC lines.

We assessed whether the hiPSC-derived sympathetic neurons had properties similar to those of their native counterparts. To our knowledge, this is the first study that assess the percentage of hiPSC-sympathetic neurons that respond to nicotine. We found that 10μM nicotine, a concentration previously reported to activate the neurons in the intact stellate ganglia preparation (Scalco *et al*., 2025), elicited increase in intracellular Ca^2+^ in 89-96% of excitable hiPSC-derived cells. Using patch clamp recordings, the same concentration of nicotine evoked membrane depolarisation and action potential generation in 84% of neurons. We also characterised the electrophysiological properties of the cells and we found that the neurons had comparable electrophysiological properties to the hiPSC-sympathetic neurons generated from the original protocol (Winbo *et al*., 2020) and to rodent sympathetic ganglia (Davis *et al*., 2020; Verkerk *et al*., 2025). They could also be grouped according to their firing patterns (tonic and phasic) as reported in other hiPSC-sympathetic neurons (Winbo *et al*., 2020; Li *et al*., 2026) and in rodent sympathetic ganglia, with our data more consistent with a majority of tonic-firing neurons described by (Luther & Birren, 2009). Other studies reported a higher percentage of phasic neurons in rat/mice (Davis *et al*., 2020; Verkerk *et al*., 2025).

Sympathetic neurons express functional adrenergic receptors, both β 2 (Bardsley *et al*., 2018) and α2 (Gilsbach *et al*., 2010; Gilsbach & Hein, 2012) subtypes. However, in control conditions, the selective β AR agonist isoprenaline results in only a modest increase in cAMP and PKA activity (Bardsley *et al*., 2018). By contrast, α2ARs have been widely reported to have an inhibitory role in sympathetic ganglia, including regulation of membrane excitability, e.g. eliciting hyperpolarisation (Brown & Caulfield, 1979), modulation of N-type Ca^2+^ channels (Lipscombe *et al*., 1989; Shapiro *et al*., 2001) and reduced neurotransmitter release (Trendelenburg *et al*., 2001; Gilsbach & Hein, 2012) via a PTx dependent pathway (Koh & Hille, 1997; Trendelenburg *et al*., 2001).

Consistent with a prevalent inhibitory role of adrenergic receptors in sympathetic neurons, norepinephrine application reduced the excitability of our hiPSC-sympathetic neurons, the effect was more pronounced when the selective α2ARs agonist (UK14304) was employed. Since norepinephrine and UK14304 are both full agonists at α2ARs, our results suggested a reduced effect when norepinephrine was employed, possibly due to its action also on β ARs therefore mitigating its effect on neuronal excitability.

We further showed that this effect was mediated, to a large extent, by GIRKs, as the inhibitory effect of the α2ARs agonist was abolished by the GIRK selective blocker Tertiapin Q. The coupling between α2ARs and GIRK has been extensively studied in recombinant systems before (Jeong & Ikeda, 1998; Bünemann *et al*., 2001; Benians *et al*., 2003; Leaney *et al*., 2004). In sympathetic neurons, most of the published works on the modulation of GIRK channels by α2ARs has been undertaken upon overexpressing GIRK subunits (GIRK1 and GIRK2 or GIRK1 and GIRK4) due to a small basal GIRK current in rat dissociated superior cervical ganglia neurons, increased upon α2ARs (Ruiz-Velasco & Ikeda, 1998; Fernández-Fernández *et al*., 2001). To our knowledge, our study is the first to demonstrate the contribution of GIRKs downstream α2ARs in human sympathetic neurons and without receptors overexpression. We validated our functional data for GIRK expression using RT-qPCR from hiPSC-sympathetic neurons and compared against published scRNAseq of the neuronal cluster of mouse thoracic sympathetic ganglia. Interestingly, our RT-qPCR data supported expression of GIRK1, GIRK2 and GIRK3 subunits which is consistent with the variable response to 1 μM ML297 reported here, indicating a possible heterogenous expression of the distinct GIRK subunit, at least in our system.

α2ARs-deficient mice are more susceptible to cardiac fibrosis, hypertrophy and heart failure as a result of cardiac pressure overload (Brede *et al*., 2002). Patients with heart failure have been reported to be less sensitive to α2ARs agonists (e.g. clonidine) possibly due to the higher levels of catecholamines reported in chronic heart disease, resulting in α2ARs desensitization (Aggarwal *et al*., 2001). Our work would therefore suggest that modulators of GIRK channels in sympathetic neurons could be a potential strategy to overcome disruption at the α2ARs level with hiPSC-sympathetic neurons as a human platform to explore this pathway.

Beyond autoregulatory mechanisms, sympathetic neuronal activity can be modulated by parasympathetic neurons. Vagal activity reduces the release of sympathetic neurotransmitters (e.g. norepinephrine and neuropeptide Y in the atria) (Koh & Hille, 1997; Jani *et al*., 2025) with muscarinic M2 receptors reported to play a central role (Trendelenburg *et al*., 2003; Jani *et al*., 2025).

Previous studies have suggested an involvement of both M1 and M2 subtypes in sympathetic ganglia. M1, via a PTx-insensitive pathway, is associated with modulation of the M current (Yang *et al*., 2006; Salzer *et al*., 2017) often studied as an isolated current rather than on neuronal excitability. By contrast, muscarinic agonists have been shown to inhibit voltage-gated Ca^2+^ channels (Shapiro *et al*., 2001; Yang *et al*., 2006) and reduce neurotransmitter release (Koh & Hille, 1997) via PTx sensitive mechanisms, with distinct contribution of M2 and M4, respectively, in mice and rat sympathetic ganglia (Shapiro *et al*., 2001). An involvement of GIRK downstream M2 muscarinic receptors via a Gα_i_-pathway has been suggested upon overexpression of GIRK1 and GIRK2 in rat sympathetic (superior cervical) ganglia (P15-21) (Fernandez-Fernandez *et al*., 1999; Fernández-Fernández *et al*., 2001).

Consistent with the published literature and available scRNAseq, we observed a high expression of *CHRM2* in hiPSC-sympathetic neurons, which was further supported by our functional data. Carbachol did not evoke an increase in intracellular Ca^2+^, and our current clamp recordings highlighted the role of muscarinic receptors in reducing sympathetic neurons excitability. Furthermore, in accordance with the coupling of M2 via Gα_i/o_, pertussis toxin abolished the effect of carbachol. Moreover, the dramatic inhibition of the carbachol effect in the presence of Tertiapin-Q strongly indicate a key involvement of GIRK channels in the muscarinic signalling pathway, previously mainly dissected in overexpression systems in rodent dissociated superior cervical ganglia (Fernandez-Fernandez *et al*., 1999; Fernández-Fernández *et al*., 2001). The contribution of other muscarinic receptors signalling pathways or the involvement of other ion channels (e.g. N-type Ca^2+^ channels, M current), which will be the scope of our future studies, might be masked by the overall inhibitory role of M2 muscarinic receptors and the prominent involvement of GIRK.

Our work therefore highlights an underappreciated role of M2 receptors in modulating sympathetic ganglia excitability beyond their well characterised function in heart rate regulation via the sinoatrial and atrioventricular regions (Ang *et al*., 2012) and ventricular contraction (Finlay *et al*., 2017). *Loci* in the *CHRM2* gene have been associated with dilated cardiomyopathy (Zhang *et al*., 2008) and reduced heart rate recovery after exercise (Ramírez *et al*., 2018), which is mediated by the autonomic nervous system and results in higher cardiovascular mortality. *Chrm2* knockout mice present elevated heart rate upon isoprenaline challenge and impaired ventricular function (LaCroix *et al*., 2008). Overall, our work could suggest that M2 receptors in sympathetic neurons may contribute to these conditions, potentially through disruption of their modulation of sympathetic excitability including under vagal tone.

We also demonstrated that the direct activation of GIRKs inhibit sympathetic neuron activity in a similar fashion, if not more pronounced, compared to the α2AR and M2 agonists, consistent with the reduced desensitization of GIRK currents upon activation with ML297 compared to GPCR agonists (Wydeven *et al*., 2014). Our data suggest a contribution of GIRK3-containing channels or heterogenous subunit composition of GIRK channels, as the lower concentration of ML297 selective for GIRK1-2 (Kaufmann *et al*., 2013) resulted in mild effects compared to the significant inhibition using 10μM. GIRK3 subunit beyond forming functional channels together with GIRK1 or GIRK2, and possibly GIRK1-GIRK2-GIRK3 (Lüscher & Slesinger, 2010), can regulate GIRK channel activity, including suppression of GIRK2-containing channels as well as trafficking, interaction with proteins involved in GPCR sensitivity (e.g. RGS2) and degradation pathways (late endosome/lysosomes) (Luo *et al*., 2022). Previous studies have reported a small GIRK-current in native rat superior cervical ganglia, with the majority of work from P15-20 rats (Fernandez-Fernandez *et al*., 1999; Fernández-Fernández *et al*., 2001). The small current could be the result of developmental regulation, with adult (P60) GIRK subunit expression and plasma localisation observed to be distinct to P15-20 in the brain (Fernández-Alacid *et al*., 2011). Another explanation is species differences, or potential differences in GIRK genes expression across sympathetic ganglia, the emerging efforts in transcriptomics analyses will enable future further examination; one of the few available superior cervical ganglia dataset (Li *et al*., 2025*a*) reported a low expression of GIRK genes and higher expression of *Chrm1* compared to *Chrm2*. Differences in protein expression and their interaction have been identified between rats and human in some other peripheral neurons (Schwaid *et al*., 2018). Little work has been undertaken on the involvement of GIRK in human neurons, with hiPSC-derived neuronal models proposed to be a strategy to identify this research gap (Luo *et al*., 2022) therefore highlighting the strength of our human *in vitro* system.

GIRK dysfunction has been described in cardiac (Gada *et al*., 2023) and neuronal disorders (Luo *et al*., 2022) and modulating their activity has been proposed as a potential therapeutic strategy (Bidaud *et al*., 2020; Luo *et al*., 2022). Due to the lack of selective modulators of GIRK3 containing channels, future efforts could focus on the development of subunit-selective GIRK activator or gene therapy approaches exploiting the distinct spatial distribution of sympathetic neurons soma and cardiomyocytes or using selective sympathetic promoters to enhance GIRK expression or activity in stellate ganglia.

In conclusion, our work supports the employment of hiPSC-sympathetic as a human *in vitro* platform for drug screening to study modulatory pathways in sympathetic neurons involved in autoregulation and parasympathetic-sympathetic crosstalk. We provide novel mechanistic insights in how adrenergic and muscarinic receptors participate in the modulation of sympathetic neurons excitability and identify GIRK channels as a converging ion channel effector in both pathways. Targeting GIRK channels in sympathetic neurons may therefore represent a novel neuromodulatory approach to reduce sympathetic drive.

## Authors contribution

L.F conceived and designed the study, performed the majority of the experiments including RT-qPCR, immunocytochemistry, whole cell patch clamp recordings, maintained and differentiated the cells, analysed the data and wrote the manuscript. M.M. designed, performed and analysed the Ca^2+^ imaging experiments. A.T. provided feedback throughout the project and assisted in the experimental design. D.A. participated in the design of the study and provided feedback throughout the project. This work was funded by grants awarded to DA. All authors contributed in the editing of the manuscript.

## Data availability

The data from this work are available from the corresponding author upon reasonable request.

## Acknowledgements

We would like to thank the Nikon Centre at King’s College London for their expertise and guidance for confocal imaging. We acknowledge King’s College London as the source of the HPSI0714i-kute_4 iPSC line, generated under the Human Induced Pluripotent Stem Cell Initiative funded by a grant from the Wellcome Trust and Medical Research Council, with support by the Wellcome Trust (WT098051) and the NIHR/Wellcome Trust Clinical Research Facility. We also acknowledge Life Science Technologies Corporation as the provider of Cytotune. The graphic abstract was created with Biorender.com

